# A propensity to hyperinsulinaemia underpins diet-induced diabetes in NOD^*k*^ mice

**DOI:** 10.64898/2026.07.19.739450

**Authors:** Matthew F Waters, Ainy Hussain, Viviane Delghingaro-Augusto, Muhammad Shamoon, Amita Bansal, Zhi-Ping Feng, T Daniel Andrews, Tenzin Dagpo, Mark E Koina, Jane E Dahlstrom, Christopher J Nolan

## Abstract

**Aims/hypothesis:** Heterogeneity in the pathophysiology of type 2 diabetes is increasingly being realised. The currently available rodent models of type 2 diabetes all have limitations and do not accurately reflect all human type 2 diabetes subtypes. NOD.BR-*H2*^*k*^ /Wicker mice (NOD^*k*^), derived from the non-obese diabetic (NOD) mouse, are type 1 diabetes resistant. However, transgene induced beta-cell stress in male NOD^*k*^ mice induces hyperinsulinaemia followed by diabetes. Here we have investigated the propensity of NOD^*k*^ mice to develop a Western-diet (WD) induced hyperinsulinaemic subtype of type 2 diabetes. Comparator mouse strains used were BALB/c and B10.BR-*H2*^*k*^ /SgSnJ mice (B10^*k*^).

**Methods:** In the longer-term studies (14-24 weeks), NOD^*k*^, B10^*k*^ and BALB/c mice were randomised to receive Chow or WD from 4 weeks of age, followed by serial measurement of body weight and fed-state blood glucose. IPGTT and IPITT tests were conducted at 13 weeks of age. Blood and pancreas were harvested for further analyses at 14 and 24 weeks of age, or sooner if diabetes developed (blood glucose concentrations ≥20 mmol/l on two consecutive days). In the acute studies, metabolic characteristics of the three strains at 8 weeks of age, continued on Chow or after a 5-day WD challenge (WDC) were assessed, along with harvesting pancreas on day 5 for *ex vivo* islet insulin secretion, electron microscopy, and bulk islet transcriptomics analyses.

**Results:** Male WD-fed NOD^k^ mice became markedly hyperinsulinaemic, gained excess weight and developed a severe type 2 diabetes phenotype. Emergence of diabetes was associated with islet endocrine cell apoptosis and loss of beta-cell mass, without evidence of insulitis. Insulin resistance on IPITT testing, however, was not evident in Chow-fed NOD^*k*^ mice. In contrast, male B10^k^ mice already had poor glucose tolerance on Chow diet and, despite having a hypoinsulinaemic phenotype, were resistant to WD-induced diabetes. BALB/c mice developed very mild glucose intolerance and hyperinsulinaemia in response to the WD. Female NOD^*k*^ mice were diabetes resistant. At 8 weeks of age, male Chow-fed NOD^*k*^ mice were mildly hyperinsulinaemic despite relative hypoglycaemia compared to the other strains. The acute 5-day WDC markedly increased hyperinsulinaemia in NOD^*k*^ mice. Transcriptomics analyses identified robust strain-specific differences, including altered islet cell differentiation, energy metabolism, endoplasmic reticulum to golgi vesicle transport and insulin processing.

**Conclusions/interpretation:** NOD^*k*^ mice, which exhibit mild hyperinsulinaemic hypoglycaemia on Chow diet and rapidly develop marked hyperinsulinaemia on WD, are type 2 diabetes prone. In contrast, B10^k^ mice have poor glucose tolerance on Chow diet and no or limited capacity to increase insulinaemia in response to WD, are diabetes resistant. These findings support the hypothesis that hyperinsulinaemia is upstream to insulin resistance in the pathogenesis of severe insulin resistant subset of type 2 diabetes for which the WD-fed NOD^*k*^ mouse is a suitable new mouse model.

**Research in Context:** *What is already known about this subject?:*

- Which of insulin hypersecretion and insulin resistance are upstream in the pathogenesis of the severe insulin resistant subtype of type 2 diabetes continues to be debated
- Rodent models of type 2 diabetes do not accurately reflect all human subtypes of type 2 diabetes
- NOD^*k*^ mice, derived from the non-obese diabetic (NOD) mouse, are type 1 diabetes resistant, but with transgene induction of islet beta-cell stress develop hyperinsulinaemia, followed by type 2 diabetes

*What is the key question?:*

- Could Western-diet fed NOD^*k*^ mice be developed as a model of severe insulin resistant type 2 diabetes and shed light on its upstream pathogenesis?

*What are the new findings?:*

- Male NOD^*k*^ mice tend to hyperinsulinaemic hypoglycaemia on Chow diet, rapidly develop marked hyperinsulinaemia on Western-diet feeding, and then develop type 2 diabetes
- Male B10^*k*^ mice (one of two comparator strains (B10^k^ and BALB/c)) have poor glucose tolerance on Chow diet, limited capacity to increase insulinaemia in response to Western-diet feeding, but are resistant to develop Western-diet induced type 2 diabetes
- Isolated islet findings show strain differences that favour intrinsic hyper-responsiveness and hypo-responsiveness of islet beta-cells of NOD^*k*^ and B10^*k*^ mice, underpinning their respective metabolic phenotypes How might this impact on clinical practice in the foreseeable future? - The findings are in support of the insulin hypersecretion hypothesis for severe insulin resistant type 2 diabetes, such that therapies to limit islet beta-cell hyperresponsiveness to prevent and treat this subtype of diabetes warrant investigation

## Introduction

While the hyperglycaemia of type 2 diabetes is generally considered to be a failure of pancreatic islet beta-cells to sustain heightened insulin secretion necessary to compensate for insulin resistance, heterogeneity in its pathogenesis is evident [1, 2]. Subtypes of type 2 diabetes are characterised according to the predominance of either insulin secretory deficiency or greater severity of insulin resistance [3, 4]. With respect to the severe insulin resistant subtype of type 2 diabetes, a debate continues as to whether insulin hypersecretion or insulin resistance is upstream in its pathogenesis [5-11]. Resolving this debate is important, as it will guide whether new approaches to the prevention and treatment of severe insulin resistant type 2 diabetes should focus on curtailing insulin secretion or reversing insulin resistance [7, 12, 13].

Consideration of type 2 diabetes heterogeneity is necessary in choosing animal models for investigating its pathogenesis and pre-clinical testing of new therapies. Currently available small animal models for the study of type 2 diabetes, highlighting advantages and limitations of each, have been recently reviewed [14]. Here, our focus is on NOD.BR-*H2*^*k*^/Wicker mice (herein NOD^*k*^ mice) and its potential role as a new mouse model of the severe insulin resistant subtype of type 2 diabetes [15, 16].

NOD^k^ mice, derived from the non-obese diabetic (NOD) mouse model of type 1 diabetes, are protected from type 1 diabetes as they carry the autoimmune diabetes resistant *H2*^*k*^ MHC haplotype [16]. Beta-cell stress induced by beta-cell specific transgenic expression of the hen egg lysozyme (HEL) protein in male NOD^*k*^ mice (NOD^*k*^ .*insHEL*) triggers diabetes, suggesting a non-immune intrinsic islet beta-cell susceptibility to failure in this model. Female NOD^*k*^.*insHEL* were resistant to diabetes development. The pathway to diabetes in male NOD^*k*^.*insHEL* is via hyperinsulinaemia [16], which prompted us to determine the effects of stressing the islet beta-cells of NOD^*k*^ mice by Western-diet (WD).

The main aim was to determine if the WD-fed NOD^*k*^ mouse could be developed as a murine model of the hyperinsulinaemic (severe insulin resistant) subtype of type 2 diabetes. Accordingly, we studied the metabolic phenotype of standard rodent chow (Chow) compared to WD feeding to 24 weeks of age in NOD^k^ mice, with B10.BR-*H2*^*k*^/SgSnJ mice (herein B10^k^ mice) and Bagg Albino (BALB/c) mice as comparator strains [17-19]. The B10^k^ mice, a C57BL/10 substrain congenic for the C57BR *H2*^*k*^ locus, was chosen as it shares the same *H2*^*k*^ MHC with NOD^*k*^ mice and has previously been used as a NOD^*k*^ comparator [15, 16, 18]. As B10^k^ mice were found to have very poor baseline glucose tolerance, BALB/c mice were also included, a strain known to have normal baseline glycaemia and to be diabetes resistant [19]. The studies predominantly focused on male mice, as female NOD^*k*^ mice were found to be resistant to WD-induced type 2 diabetes. As we found that WD-feeding of male NOD^*k*^ mice for 4 weeks already resulted in marked hyperinsulinaemia, we undertook an additional series of short 5-day WD challenge (WDC) studies to ascertain early events in its phenotypic development, including at the level of the pancreatic islet.

## Methods

### Mice

All mice experiments were approved by the Animal Experimentation Ethics Committee of the Australian National University (ANU) (protocols: A2014/24, A2017/25, A2020/43). NOD.BR-*H2*^*k*^ /Wicker (NOD^*k*^) mice were sourced from the the Australian Phenomics Facility, ANU (Acton, ACT, Australia). B10.BR-*H2*^*k*^ /SgSnJ (B10^*k*^) mice and BALB/c mice were sourced from the Animal Resource Centre (Canning Vale, WA, Australia). Mice were housed at The Canberra Hospital Animal Facility (Garran, ACT, Australia) for the longer-term studies and at the Australian Phenomics Facility (ANU) for the 5-day WDC studies. Mice were housed in a humidity-controlled environment on a 12 h light/dark cycle with free access to water and were fed ad-libitum.

### Experimental diets

Two series of studies were performed. Mice were fed Chow diet (Gordon’s Specialty Stockfeed, Yanderra, NSW, Australia) until the commencement of experiments.

The first series involved feeding of Chow or Western-diet (WD) (SF03-020, Semi-Pure Rodent Diet, Specialty Feeds, Glen Forrest, WA, Australia) from 4 weeks of age in 2 cohorts: (i) to 24 weeks of age or diabetes diagnosis (plasma glucose ≥20 mmol/l for 2 consecutive days) in male mice only; and (ii) to 14 weeks of age (male and female mice). Body weight and 9am tail blood glucose (StatStrip XpressTM, Nova Biomedical, Flintshire, United Kingdom) and plasma insulin were measured fortnightly. In the first cohort an IPITT and in the second cohort an IPGTT were performed at 13 weeks of age. On the last experimental day (both cohorts), a 9am fed-state tail blood sample (60 μl) was collected followed by euthanising the mice by cervical dislocation for cardiac blood sampling and pancreas harvest. Blood samples were collected into 0.5 M EDTA-rinsed microfuge tubes, stored on ice prior to centrifugation and plasma separation, followed by storage at −80°C.

The second series, the 5-day WDC experiments, involved assigning 51 (± 3 days) day old male to either Chow or WD feeding for 5 days, such that they were 8 weeks of age at study end. On day 0 and day 5 fed-state body weight, tail blood glucose (glucose meter) and 30 μl blood samples were collected for plasma insulin measurement. On day 5, mice were assigned to either a terminal intraperitoneal glucose tolerance test experiment or to be ethically culled for islet isolation for static insulin secretion, electron microscopy or bulk islet RNA sequencing studies.

On commencement of all experiments, alternate mice were strictly allocated to Chow or WD.

### ipGTT and ipITT tests

IPGTTs (1g/kg 25% glucose) and IPITTs (0.75U/kg regular insulin) were performed in fasted conscious mice, as described in the Electronic Supplemental Material (ESM).

### Plasma chemistry

Plasma insulin was measured by radioimmunoassay (Linco Research) and plasma proinsulin and C-peptide by ELISA kits (Alpco). Serum triglycerides and NEFA concentrations were measured by colorimetric enzyme kit assays (Wako Pure Chemical Industries Ltd.) and plasma adiponectin by a mouse adiponectin ELISA kit (Abcam). Potential interference in the insulin assay by insulin autoantibodies (IAA) was assessed by a non-competitive ELISA method [20, 21], and by plasma insulin assay after polyethylene glycol (PEG) (Sigma-Aldrich) precipitation, as described in the ESM [22].

### Pancreas histology, insulin immunohistochemistry and measurement of beta-cell mass

After terminal cardiac blood sampling, pancreases were quickly excised, removed of fat, weighed and fixed in 10% neutral buffered formalin solution before embedding in paraffin. Histological analysis of pancreas was conducted on 4 µm sections stained with H&E. For measurement of islet beta-cell mass, 3 non-consecutive sections of 5 μm, 100 μm apart were cut for immunostaining for insulin and analysis, as described in the ESM. Blinded analyses were performed.

### Islet isolation and static insulin secretion

Islets were isolated by Liberase digestion (Roche Diagnostics, Castle Hill, Australia), gradient centrifugation (Histopaque, Sigma-Aldrich) and handpicked under a stereomicroscope. Assessment of ex-vivo insulin secretion with islet incubation at glucose concentrations of 3 mmol/l (3G), 8 mmol/l (8G), 16 mmol/l (16G) and 8G with 100 μM isobutylmethylxanthine (IBMX) (Sigma-Aldrich) for 45 min, was performed as described in the ESM. The insulin ultrasensitive kit (Cisbio) was used to measure insulin concentrations in both the secretion media and acid-alcohol islet lysates.

### Islet electron microscopy

Freshly isolated islets were prepared for transmission electron microscopy (EM) analysis, as described in the ESM. Mature and immature insulin granules were identified based on core density and presence or absence of the characteristic mature granule halo. Blinded analyses were performed.

### RNA sequencing

Islet RNA isolation was performed using the RNeasy minikit (Qiagen). RNA concentration and integrity were assessed using the Agilent 2100 Bioanalyzer (Agilent) and the Agilent RNA 6000 Nano kit (Agilent). RNA samples with RNA integrity number >7 were used for gene expression analyses. RNA sequencing libraries were prepared using Illumina TruSeq Stranded mRNA library kit and sequenced using the Illumina NextSeq 500 system using the 75 cycle high output run kit (Illumina). Following quality check and excluding lowly expressed genes, Limma-voom was used to identify differentially expressed genes (DEGS) using normalised reads for group-wise comparisons with log_2_ Fold Change (FC), average expression across all samples in log_2_ Counts Per Million (CPM; AveExpr), p-values, and q-values after Benjamin-Hockberg false discovery rate correction. DEGs with q<0.05 and positive log_2_CPM in group comparisons were assessed via Enrichr for gene ontology enrichment to identify pathways of biological significance. To filter beta-cell specific genes from our islet bulk RNA-seq data, we interrogated with the 60-day-old murine beta-cell gene expression atlas available at EMBL-EBI (https://www.ebi.ac.uk/gxa/experiments/E-MTAB-2266/Results), as well as the mouse MSigDB HALLMARK_PANCREAS_BETA_CELLS gene set (https://www.gsea-msigdb.org/gsea/msigdb/mouse/collection_details.jsp#MH).

Gene expression was validated using qRT-PCR. RNA (200 ng) was reverse transcribed to cDNA using the QuantiTect Reverse Transcription Kit (Qiagen) according to manufacturer’s instructions. PCR was performed in a 384-well plate on the 7900 HT Real Time PCR System (Applied Biosystems). Target gene expression was normalised to housekeeping genes beta actin (*Actb*) and glyceraldehyde-3-phosphate dehydrogenase (*Gapdh*) and expressed as a fold-change in all mice groups compared to the BALB/c Chow mice group.

### Statistical analyses

For statistical analyses R (v4.3.1), RStudio (v2023.03.0+386), and GraphPad Prism (v10.0.3) were used. Results are presented as mean ± SEM or median (for ordinal data). All time-course data were tested using linear regression mixed models. Diabetes free survival was assessed using a Cox regression model. Other group comparisons were by two-way ANOVA, ordinal logistic regression, or the Kruskal-Wallis rank sum test, with *post-hoc* testing for multiple comparisons as indicated. Statistical significance was set at *p*<0.05.

## Results

### Male NOD^*k*^ mice only are prone to WD-induced hyperinsulinaemia and diabetes

Key findings were that feeding WD to male NOD^k^ mice resulted in fed-state hyperglycaemia by 10 weeks of age, glucose intolerance despite severe hyperinsulinaemia by 13 weeks of age, and the development of type 2 diabetes between 13 and 24 weeks of age (Fig. 1a-g). Feeding WD to female NOD^*k*^ mice led to the development of glucose intolerance with moderate hyperinsulinaemia by 13 weeks of age, while fed-state glucose remained minimally affected (ESM Fig. 1a-c). In contrast, male WD-fed B10^*k*^ mice were resistant to the development of diabetes (Fig 1.b,c), despite Chow and WD-fed B10^*k*^ having severe glucose intolerance and B10^*k*^ mice failing to increase insulinaemia with WD feeding (Fig. 1b-g). Male BALB/c mice were normally glucose tolerant on Chow diet with minimal change with WD feeding associated with a mild increase in insulinaemia (Fig. 1b-g). Four-week-old male NOD^*k*^ mice had the highest baseline body weight and in response to WD feeding, showed accelerated weight gain after only 2 weeks on diet, in contrast to B10^*k*^ and BALB/c mice in which accelerated weight gain on WD was not evident until after 6 weeks (Fig. 1a). IPITTs in 13 week old Chow-fed mice, showed that male NOD^*k*^ mice are insulin sensitive prior to WD feeding (Fig. 1h).

**Fig. 1.**
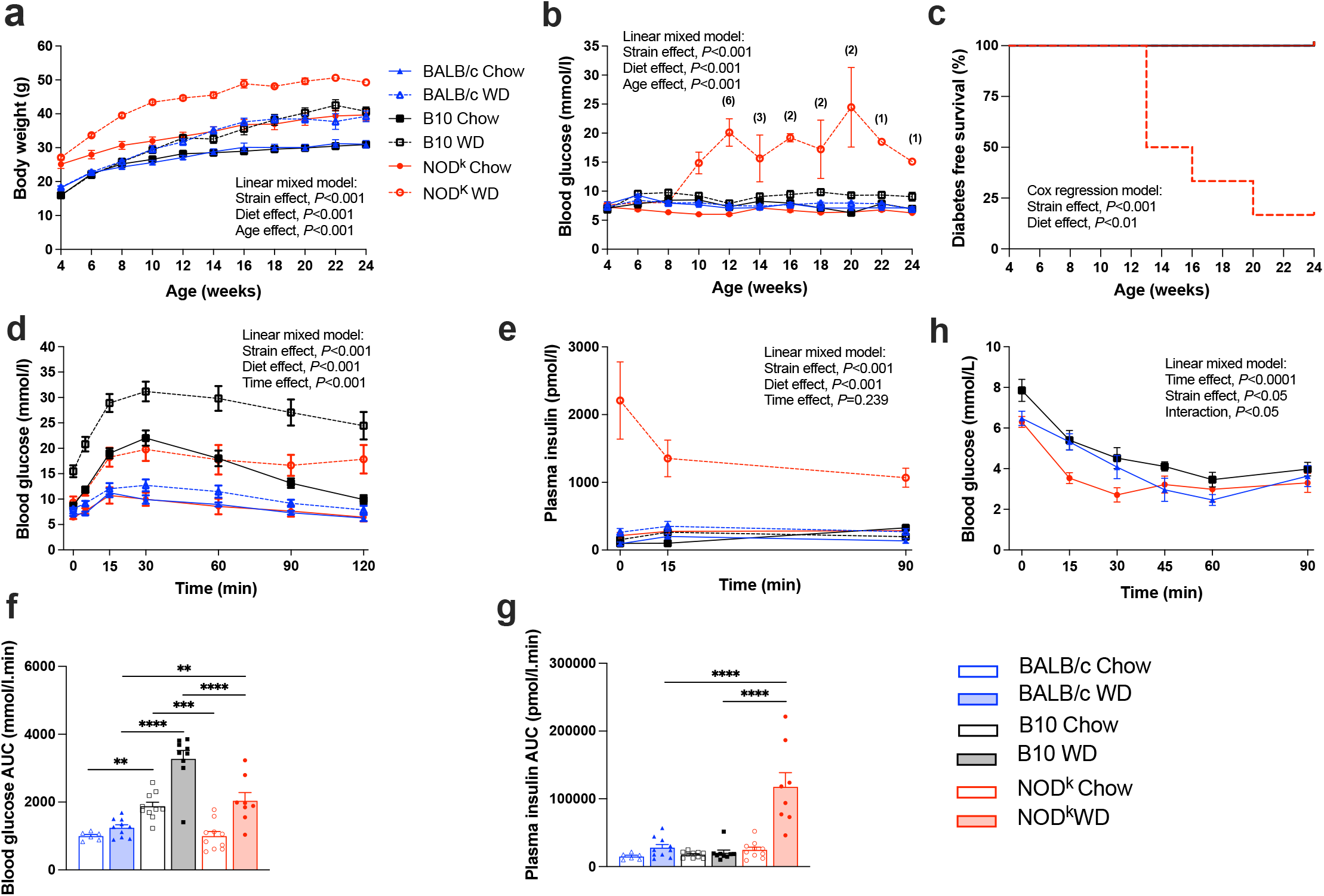
Metabolic characteristics of male BALB/c, B10^*k*^ and NOD^*k*^ mice fed Chow or Western-diet (WD) from 4 to 24 weeks of age. **(a)** Body weight, **(b)** 9am fed blood glucose, **(c)** Diabetes free survival; *n*=6 all groups; numbers in parentheses indicate NOD^*k*^ mice not yet culled because of diabetes **(b). (d-g)** IPGTTs performed at 13 weeks of age; blood glucose (d), Plasma insulin **(e)**, IPGTT blood glucose area under the curve (AUC) **(f)**, IPGTT plasma insulin AUC **(g)**; *n*=6 BALB/c Chow, *n*=10 BALB/c WD, *n*=10 B10^*k*^ Chow, *n*=9 B10^*k*^ WD, *n*=10 NOD^*k*^ Chow, *n*=8 NOD^*k*^ WD. **(h)** IPITTs performed at 13 weeks of age; *n*=6 BALB/c Chow (2 mice culled at 60 min time point because of severe hypoglycaemia), *n*=5 B10^k^ Chow, *n*=6 NOD^*k*^ Chow (1 mouse culled at 30 min due to severe hypoglycaemia). Data are shown as means ± SEM. Analysed with linear mixed model with Tukey’s *post-hoc* tests **(a, b, d, e, h)**, BALB/c Chow vs B10^k^ Chow n.s., NOD^k^ Chow vs B10^k^ Chow *p*<0.0001, NOD^k^ Chow vs BALB/c Chow *p*<0.0001, BALB/c WD vs B10^k^ WD n.s., NOD^k^ WD vs B10^k^ WD *p*<0.0001, NOD^k^ WD vs BALB/c WD *p*<0.0001 **(a)**, BALB/c WD vs B10^*k*^ WD n.s., NOD^*k*^ WD vs B10^*k*^ WD *p*<0.0001, NOD^*k*^ vs BALB/c *p*<0.0001 **(b)**, BALB/c Chow vs B10^*k*^ *p*<0.01 Chow, NOD^*k*^ Chow vs B10^*k*^ Chow *p*<0.01, NOD^*k*^ Chow vs BALB/c Chow n.s., BALB/c WD vs B10^*k*^ WD *p*<0.0001, NOD^*k*^ WD vs B10^*k*^ WD *p*<0.01, NOD^*k*^ WD vs BALB/c WD *p*<0.0001 **(d)**, BALB/c WD vs B10^*k*^ WD n.s., NOD^*k*^ WD vs B10^*k*^ WD *p*<0.0001, NOD^*k*^ WD vs BALB/c WD, *p*<0.0001 **(e)**; Cox-regression model **(c)**; and by two-way ANOVA with Tukey’s multiple comparison *post-hoc* tests **(f, g)**, \**p*<0.05, \*\**p*<0.01, \*\*\**p*<0.001, \*\*\*\**p*<0.0001.

### Male NOD^*k*^ mice are profoundly hyperinsulinaemic in response to WD-feeding by 14 weeks of age

Fed blood chemistry (9am) in 14-weeks of age Chow-fed mice showed moderately elevated plasma insulin and proinsulin in NOD^*k*^ mice compared to the other two strains, however, C-peptide was only elevated in NOD^*k*^ compared to B10^*k*^ mice (Fig. 2a-c). WD feeding to 14 weeks of age in NOD^*k*^ mice caused a marked 7.2 fold increase in insulin without any increase in C-peptide levels, suggestive of markedly reduced insulin clearance in these mice (Fig. 2a-c). The possibility of IAA interference in the insulin assay to explain these very high insulin concentrations was excluded (ESM Fig. 2). Fed-state C-peptide levels were low in both Chow and WD fed B10^k^ mice, consistent with poor insulin secretion that is unresponsive to WD-feeding in these mice.

**Fig. 2.**
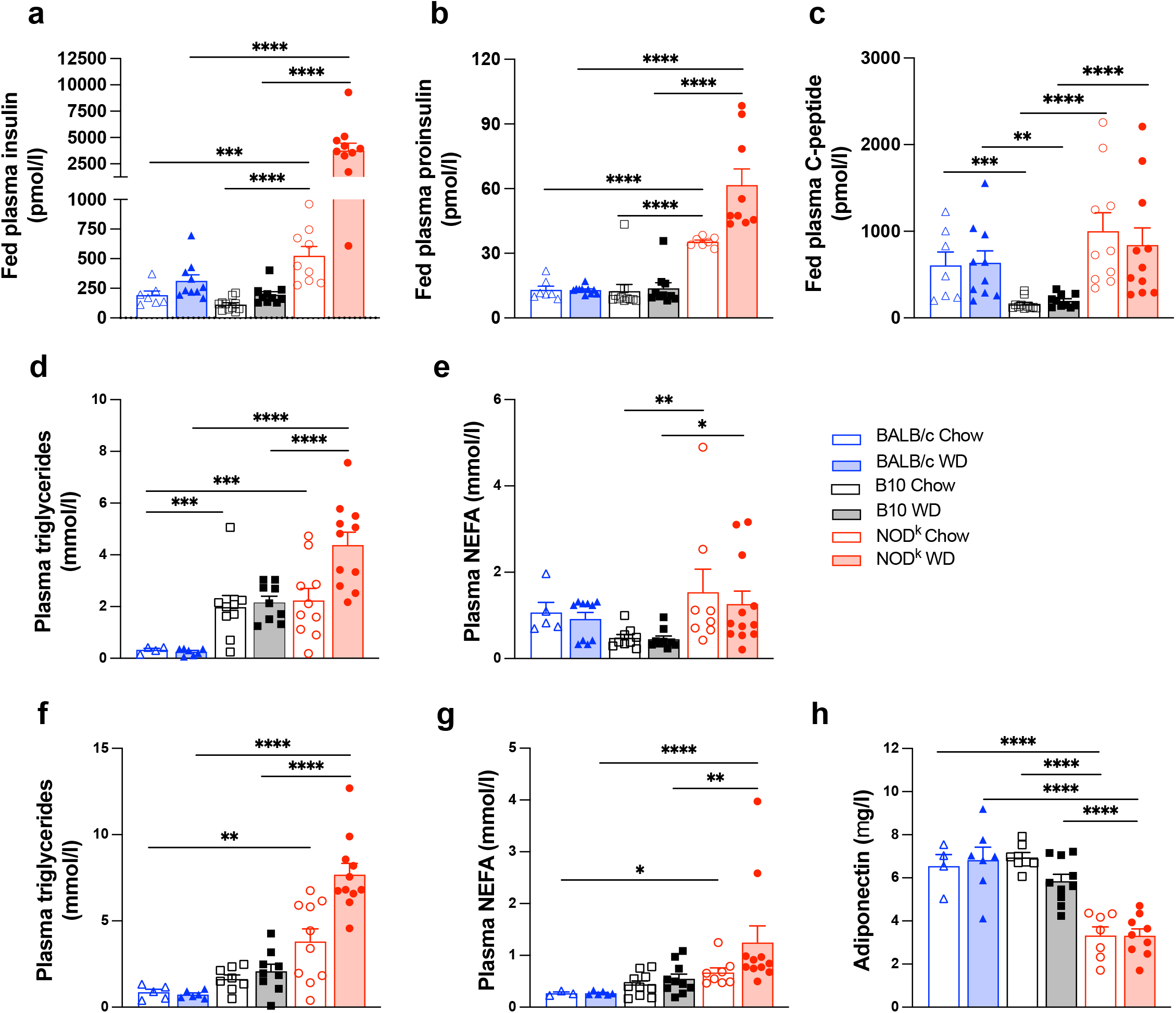
Fed plasma insulin, proinsulin, C-peptide, triglycerides, non-esterified fatty acid (NEFA) and adiponectin at 14 weeks of age and fasted triglycerides and NEFA at 13 weeks of age in male BALB/c, B10^*k*^ and NOD^*k*^ mice fed Chow and WD. (a-c) Fed plasma insulin **(a)**, Proinsulin *n*= **(b)**, and C-peptide **(c)**; *n*=7-11. **(d, e)** Fasted plasma triglycerides (d) and NEFA (e); *n*=4-12. **(f, g)** Fed plasma triglycerides **(f)** and NEFA **(g)**; n=3-11. **(h)** Fed plasma adiponectin; *n*=4-10. Data are shown as means ± SEM. Analysed by two-way ANOVA, with Tukey’s multiple comparison *post-hoc* tests, \**p*<0.05, \*\**p*<0.01, \*\*\**p*<0.001, \*\*\*\**p*<0.0001.

### Male NOD^*k*^ have unfavourable lipid and adiponectin profiles

Male NOD^*k*^ mice were hypertriglyceridaemic in both the fasted and fed-state compared to BALB/c mice at 13-14 weeks of age, evident in Chow-fed, but worsened in WD-fed mice (Fig. 2d, f); whereas NEFA levels were only elevated in NOD^*k*^ compared to BALB/c mice in the fed-state (Fig. 2e, g). B10^*k*^ mice had evidence of fasting hypertriglyceridaemia compared to BALB/c mice, without evidence of WD worsening or elevated NEFA (Fig. 2d-g). Male NOD^*k*^ mice, Chow and WD fed, had markedly reduced fed-state plasma adiponectin levels compared to BALB/c and B10^*k*^ mice (Fig. 2h).

### Exocrine tissue lipid and pancreas inflammation

Male NOD^*k*^ mice pancreases showed moderate lipid droplet accumulation in WD-fed mice by 14 weeks of age (ESM Fig. 3a-c), with moderate accumulation in older Chow and WD-fed mice (Fig. 3a-c). Of note, WD-fed BALB/c mice also accumulated exocrine tissue lipid droplets, evident at 14 weeks and increased by 24 weeks of age (ESM Fig. 3a-c; Fig. 3a-c). B10^*k*^ mice pancreases did not accumulate lipid droplets. A mild to moderate degree of pancreatic periductal inflammation was present in NOD^*k*^ and B10^*k*^, without evidence of worsening with WD-feeding (Fig. 3b, d). Inflammation was mostly absent in BALB/c pancreases. Peri-islet inflammation was always in association with periductal inflammation (Fig. 3b, d). Invasive insulitis was not seen in any mice groups.

**Fig. 3.**
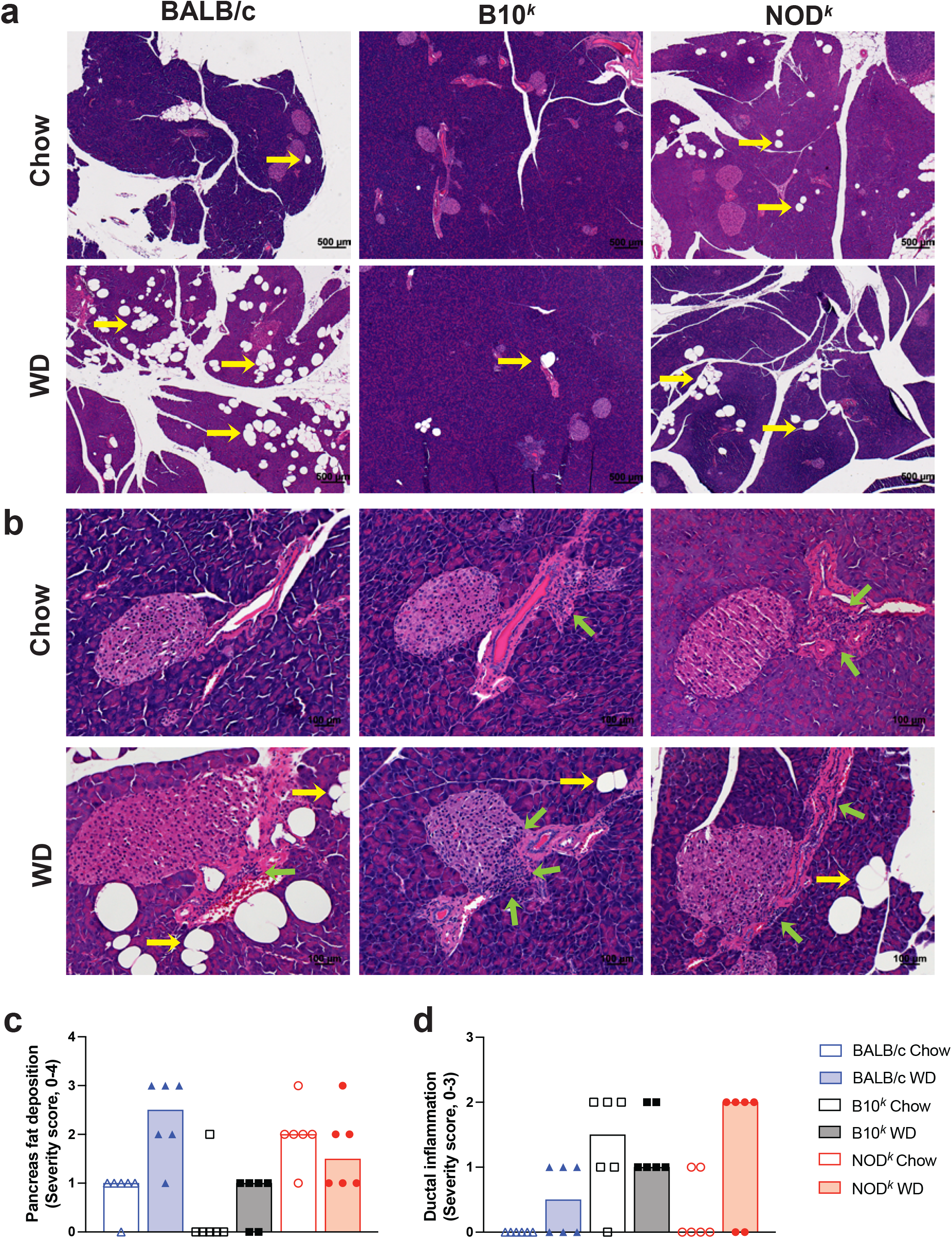
Pancreas histology in male BALB/c, B10^*k*^ and NOD^*k*^ mice fed Chow and WD to 24 weeks of age. **(a)** Representative low power images of pancreatic sections stained with H&E showing presence of fat deposits; scale bar 500 μm, initial magnification x50; yellow arrows point to fat deposits. **(b)** Representative high power images of pancreas sections stained with H&E showing periductal inflammation; scale bar 50 μm, initial magnification x400; green arrows point to areas of inflammation. **(c)** Pancreatic fat deposition scores; score matrix 0-none, 1-minimal, 2-mild, 3-moderate, 4-severe. **(d)** Periductal inflammation scores; score matrix 0-none, 1-<10%, 2-10-50%, 3->50% of ducts affected. Each data point represents one mouse and the bars are the group median. Ordinal logistic regression model **(c**,**d)**; diet effect, *p*=0.0148, strain effect, *p*=0.0009, diet × strain effect, *p*=0.0157 **(c)**; diet effect, *p*=0.0323, strain effect, *p*=0.0067 **(d)**.

### WD-fed male NOD^*k*^ islets have more apoptotic endocrine cells and lower beta-cell mass as diabetes develops

While islet beta-cell mass is maintained in male WD-fed NOD^*k*^ mice by 14 weeks of age, beta-cell mass loss is evident in these mice at the later age as diabetes develops (ESM Fig. 4a, b; Fig. 4b-d). Islet endocrine cell apoptosis was more commonly seen in older WD-fed NOD^*k*^ islets (Fig. 4a), and IHC insulin immunostaining was more patchy and less intense in these mice (Fig. 4b) indicative of beta-cell dysfunction. Additionally intracytoplasmic inclusion bodies were commonly seen in the endocrine cells of the older WD-fed NOD^*k*^ mice, but not in BALB/c and B10^*k*^ mice (Fig. 4a).

**Fig. 4.**
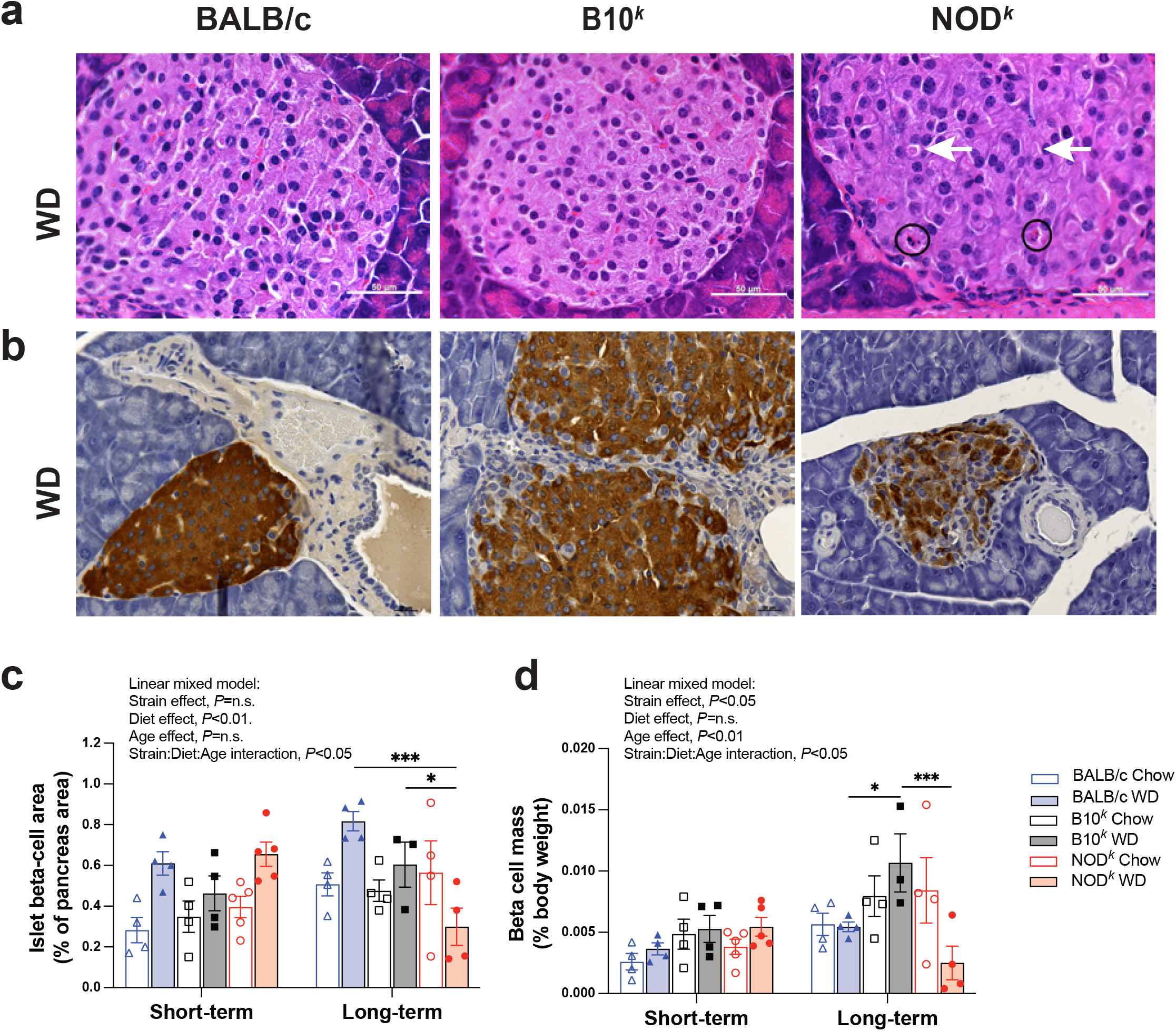
Islet cell apoptosis, insulin immunohistochemistry and islet beta-cell mass assessment in male BALB/c, B10^*k*^ and NOD^*k*^ mice fed Chow and WD. **(a)** Representative high power images of pancreas sections of 24 week old mice stained with H&E showing apoptosis and intracytoplasmic inclusion bodies. In these images apoptotic endocrine cells, as shown by the circling of apoptotic cells, and endocrine cell intracytoplasmic inclusion bodies, as shown by the white arrows, are evident in the WD-fed NOD^k^ mouse only; scale bar 50 μm, initial magnification x630. **(b)** Representative high power images of pancreas sections of 24 week old mice immunostained for insulin (brown). Reduced intensity of insulin staining is evident in the WD-fed NOD^*k*^ mouse only; scale bar 50 μm, initial magnification x400. **(c, d)** Islet beta-cell area (as percentage of pancreas area) **(c)** and islet beta-cell mass (as a percentage of body weight) in mice harvested at 14 weeks of age (short-term) and 24 weeks of age (or earlier if culled because of diabetes, long-term); *n*=4 BALB/c both diets, *n*=5 NOD short term and *n*=4 NOD^k^ long term bother diets, *n=4* B10^k^ short term both diets and long term Chow diet, *n*=3 B10^*k*^ long term WD. Data are shown as means ± SEM. Analysed by linear mixed model with Tukey’s multiple comparison *post-hoc* tests, \*\**p*<0.01, \*\*\**p*<0.001.

### Young male NOD^*k*^ mice display a hyperinsulinaemic hypoglycaemic phenotype which is accentuated by 5-days of WD feeding

To investigate the early effects of nutrient-induced stress on the metabolic phenotypes of the three mice strains, we next performed a 5-day WDC study in 8-week-old male mice. Key findings were: (i) relative hypoglycaemia of Chow-fed NOD^*k*^ mice compared to the other two strains during IPGTT testing, as well as in the fed state, which was not altered by 5-days of WD feeding (Fig. 5c, e, g); and (ii) the WDC markedly accentuated the IPGTT and fed-state hyperinsulinaemia of NOD^k^ mice (Fig. 5d, f, h). Chow-fed NOD^k^ mice were again found to be heavier than BALB/c and B10^*k*^ mice (Fig. 5a), although the percentage body weight gain was similarly accelerated by the WDC in BALB/c and NOD^*k*^, but not in B10^*k*^ mice (Fig. 5b).

**Fig. 5.**
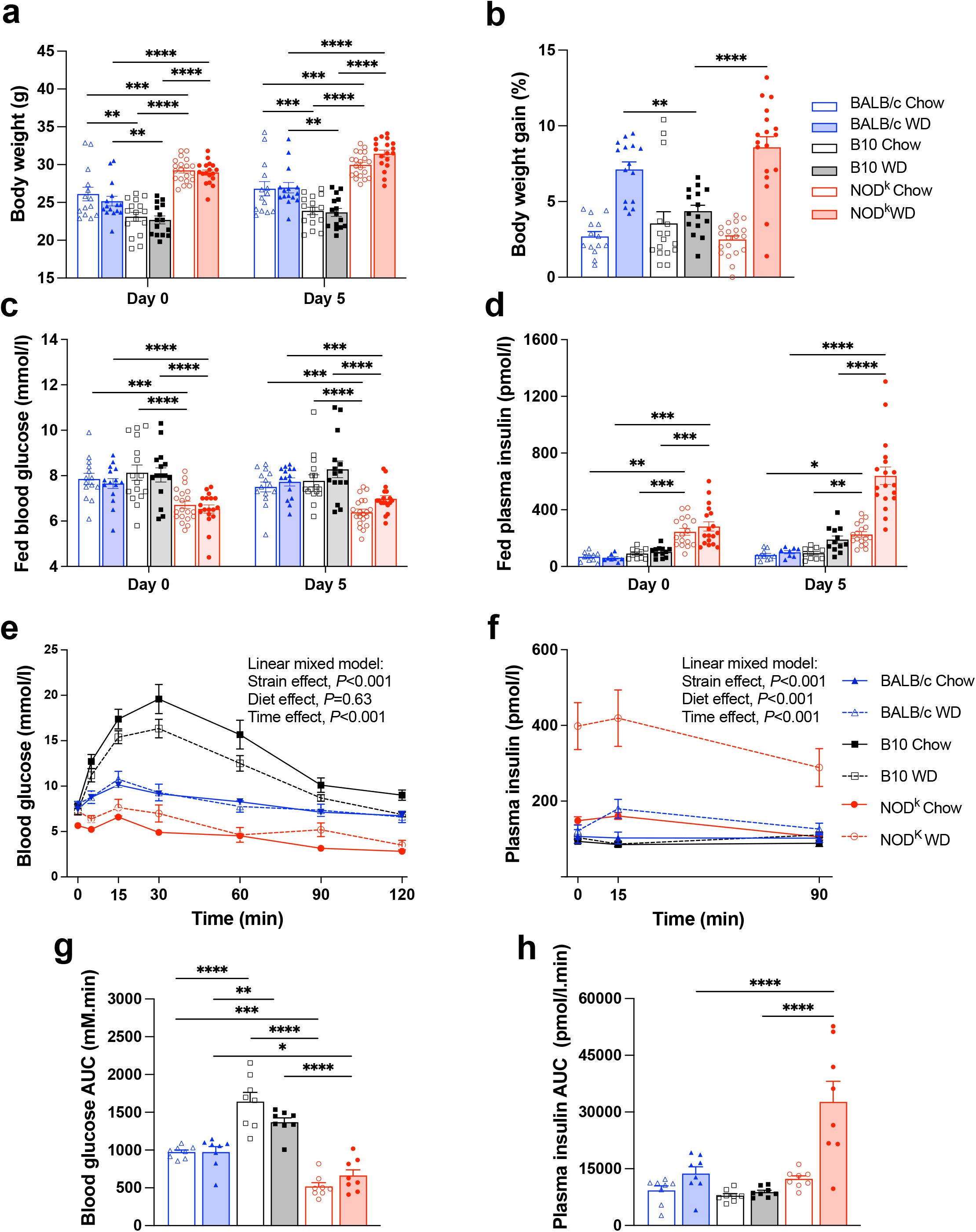
Body weight, fed blood chemistry and IPGTT results of 8 week old BALB/c, B10^*k*^ and NOD^*k*^ mice continued either on Chow diet or in response to a 5-day WDC. **(a-d)** Body weight (days 0 and 5) **(a)**, Percentage change in body weight day from day 0 to 5 **(b)**, Fed 9am blood glucose (days 0 and 5) **(c)**, Fed 9am plasma insulin (days 0 and 5) **(d)**; *n*=14 BALB/c Chow, *n*=15 BALB/c WD, *n*=16 B10^*k*^ Chow, *n*=15 B10^*k*^ WD, *n*=20 NOD^*k*^ Chow, *n*=18 NOD^*k*^ WD **(a, b, c)**, *n*=10 BALB/c Chow, *n*=12 BALB/c WD, *n*=10 B10^*k*^ Chow, *n*=8 B10^*k*^ WD, *n*=17 NOD^*k*^ Chow, *n*=18 NOD^*k*^ WD **(d). (e-h)** IPGTTs performed at day 5; Blood glucose **(e)**, Plasma insulin **(f)**, IPGTT blood glucose AUC **(g)**, IPGTT plasma insulin AUC **(h)**; *n*=8 for all groups. Data are shown as means ± SEM. Analysed with linear mixed model with Tukey’s *post-hoc* tests **(a, c-f)**; BALB/c Chow vs B10^k^ Chow *p*<0.0001, NOD^k^ Chow vs B10^k^ Chow *p*<0.0001, NOD^k^ Chow vs BALB/c Chow *p*<0.001, BALB/c WD vs B10^*k*^ WD *p*<0.01, NOD^*k*^ WD vs B10^*k*^ WD *p*<0.0001, NOD^*k*^ WD vs BALB/c WD *p*<0.01 **(e)**, BALB/c WD vs B10^*k*^ WD n.s., NOD^*k*^ WD vs B10^*k*^ WD *p*<0.0001, NOD^*k*^ WD vs BALB/c WD *p*<0.0001 **(f)**; and by two-way ANOVA with Tukey’s multiple comparison *post-hoc* tests **(b, g, h)**, \**p*<0.05, \*\**p*<0.01, \*\*\**p*<0.001, \*\*\*\**p*<0.0001.

### Glucose-stimulated insulin secretion (GSIS) in *ex vivo* islets of 8 weeks of age male mice

The 5-day WDC compared to Chow diet had no significant effects on *ex vivo* islet insulin content or insulin secretion in any of the mice groups. In comparing GSIS of the three strains, and expressed as a percentage of insulin content, insulin secretion in response to 8G+IBMX was mildly increased in islets of NOD^*k*^ mice compared to those of the other two strains (Fig. 6a). However, when expressed as the fold change from 3G Chow-fed mice islets of the same strain, insulin secretion at 16G and 8G+IBMX was increased in NOD^*k*^ by approximately 60% to 130% compared to BALB/c and B10^*k*^ islets (Fig. 6b).

**Fig. 6.**
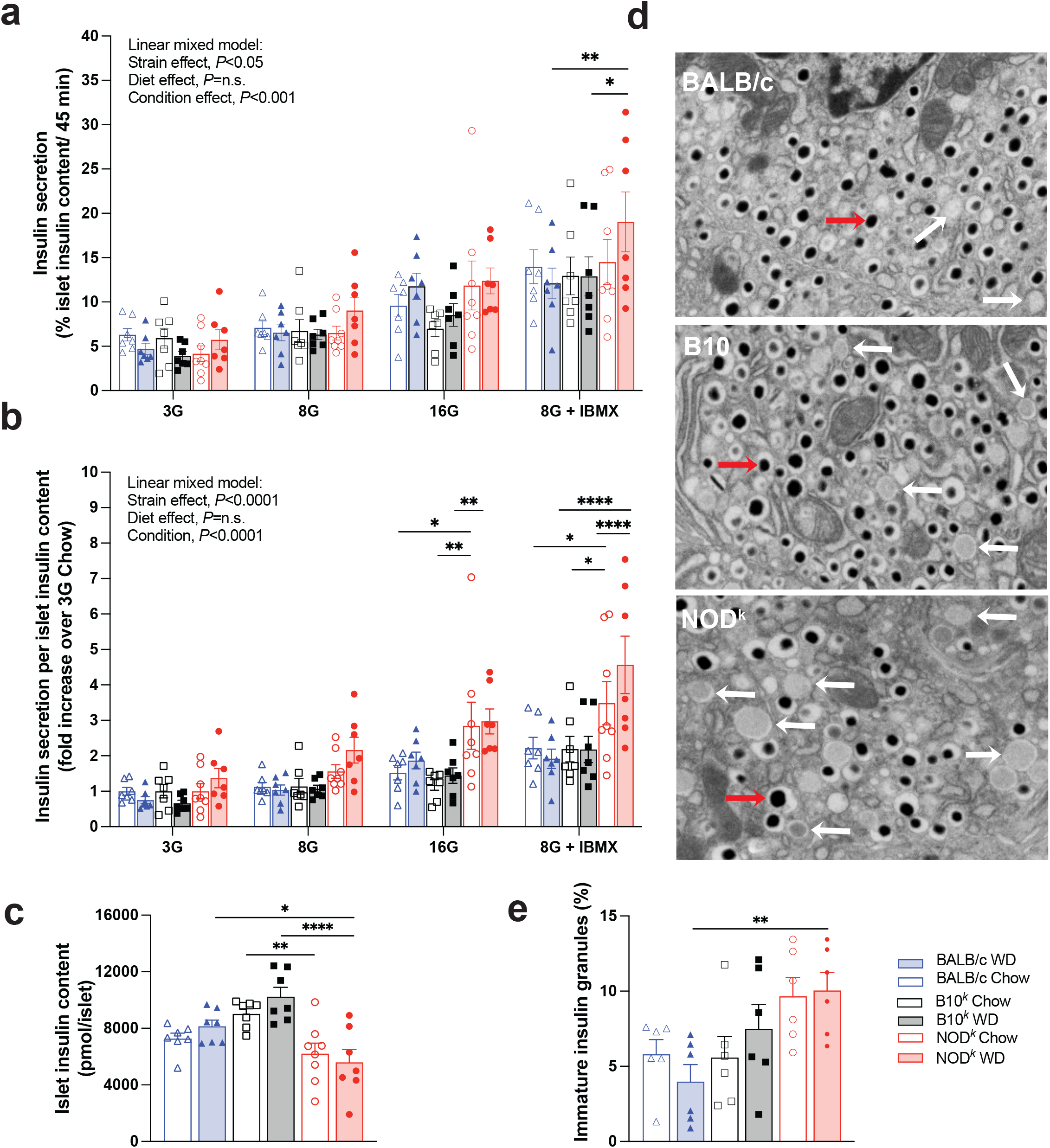
*Ex vivo* islet insulin secretion, islet insulin content and transmission electron microscopy analysis of isolated islet ultrastructure. **(a-c)** Static insulin secretion measured in islets, isolated from 8-week-old male BALB/c, B10^k^ and NOD^k^ mice continued either on Chow diet or in response to a 5-day WDC incubated at 3 (3G), 8 (8G) and 16 (16G) mmol/l glucose, and 8G + 100 μM IBMX. Results expressed as the percentage of islet insulin content secreted over 45 min **(a)**, and as the fold change in insulin secretion from 3G Chow-fed mice islets of the same strain normalised by total insulin content **(b)**, and islet insulin content **(c)**; each data point is from the average of triplicate wells; *n*=*7* mice in all groups, except *n*=8 for the NOD^k^ Chow group. **(d, e)** Representative electron photomicrographs of islet beta-cells from male WD-fed BALB/c, B10^*k*^ and NOD^*k*^ mice; red arrows point to mature insulin granules and white arrows point to immature insulin granules **(d)**, and percentage of immature insulin granules **(e)**; each data point is from the average of 10-12 islet beta-cells from 5 islets per mouse; *n*=6 mice per group. Data are shown as means ± SEM. Analysed with linear mixed model with Tukey’s *post-hoc* tests **(a, b)**; and by two-way ANOVA, with Tukey’s multiple comparison *post-hoc* tests **(c, d)**, \**p*<0.05, \*\**p*<0.01, \*\*\*\**p*<0.0001.

### Islet beta-cell ultrastructure reveals an increased percentage of immature insulin granules in NOD^k^ mice

Overall, no differences between strains were seen in the total number of insulin granules per islet beta-cell area (results not shown). However, when comparing WD-fed mice, NOD^*k*^ showed a higher proportion of immature insulin granules compared to BALB/c mice (Fig. 6d, e). The percentage of immature granules within B10^*k*^ mice islets was not different to either of the other mouse strains (Fig. 6e).

### Islet transcriptomic profiles show clear strain differences between mice strains

Bulk transcriptomic profiles of islets were assessed using unbiased RNA-seq in 8-week-old male mice continued on Chow diet or challenged by 5 days of WD feeding. The islet transcriptomes of the strains were clearly different, however the 5 day WD had minimal impact on gene expression within the strains (Fig.7a-c). The top DEGs between strains in this unbiased analysis are shown in Fig. 7d-f (volcano plots) and ESM Fig.4 (heat map). Of interest, NOD^*k*^ islets had higher expression of *Ckb, Slco1a6* and *Padi2* and lower expression of *Akr1e1* and *Cttnbp2* compared to BALB/c and B10^*k*^ mice; whereas B10^*k*^ mice had higher expression of *C8b* and lower expression of *Scg5* and *Scn1b* compared to NOD^*k*^ and BALB/c islets. The expression level of 6 DEGs (*Acadl, Adc5y, Pgm2l1, Rasgrp, Slc39a8* and *Tcf7l1*) identified via islet bulk RNA-seq, which were consistently highly differentially expressed in NOD^*k*^ compared to both BALB/c and B10^*k*^ mice and involved in biological pathways of significance in islet biology, was validated using qRT-PCR (ESM Fig. 6).

**Fig. 7.**
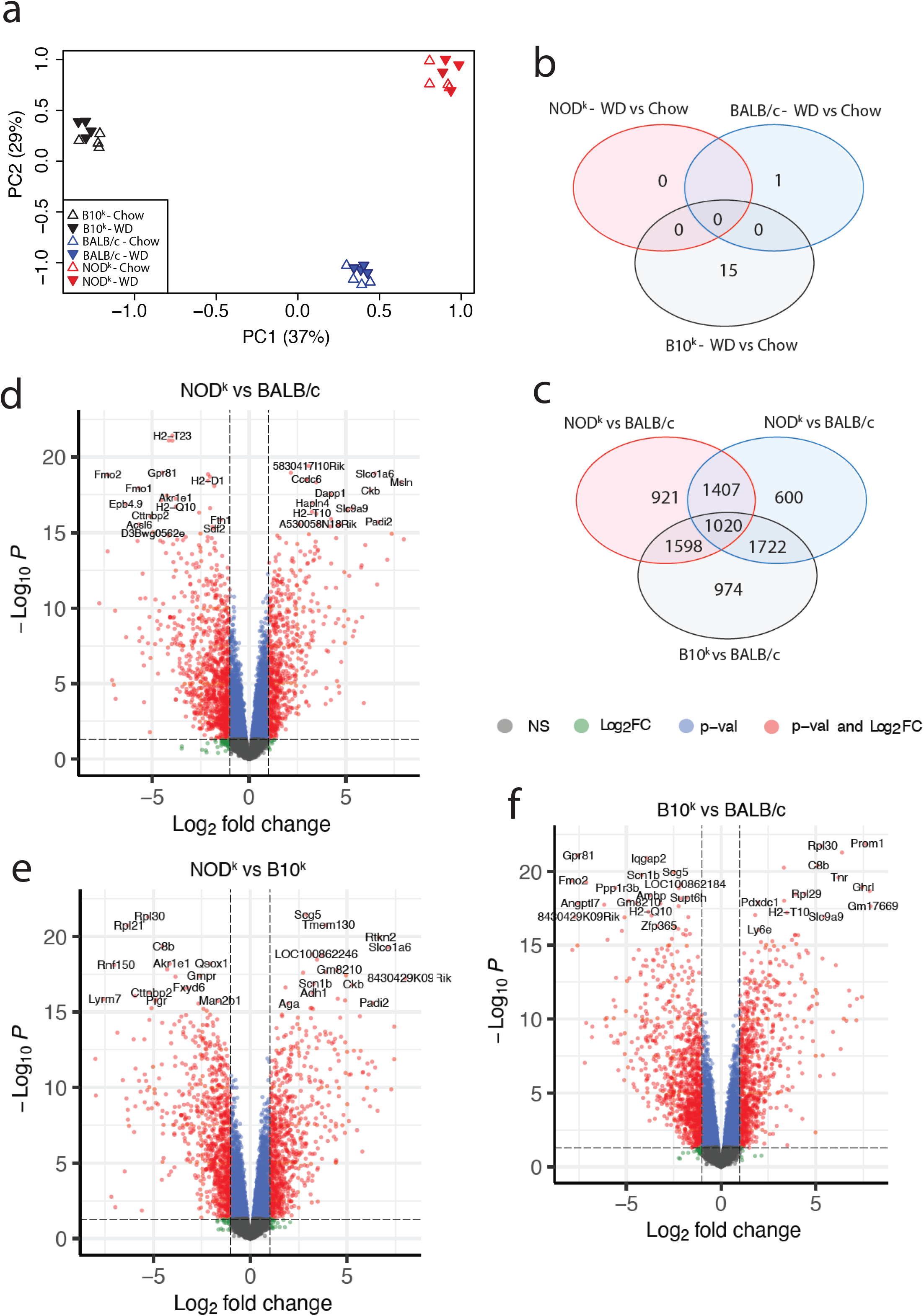
Bulk pancreatic islet transcriptome analyses of 8-week-old male BALB/c, B10^*k*^ and NOD^*k*^ mice continued either on Chow diet or in response to a 5-day WDC; *n=*4 all groups. **(a)** Principal components analysis showing transcriptome separation by strain but not diet. **(b**,**c)** Ven diagrams showing the number of differentially expressed genes (DEGs) between Chow and WD diet within strains **(b)**, and between strains (Chow and WD-fed mice of each strain combined) **(c). (d-f)** Volcano plot of DEGs between NOD^k^ and BALB/c mice **(d)**, NOD^k^ and B10^k^ mice **(e)**, and B10^k^ and BALB/c mice **(f)**. Chow and WD-fed mice within each strain combined; dashed line significance threshold *p*<0.05, log fold-change threshold >1.0 (d-f).

On filtering to the MSigDB HALLMARK_PANCREAS_BETA_CELLS gene set, the expression patterns of established islet genes differed between strains (Fig. 8a-d). Higher expression of *Dcx* and *Sst* and lower expression of *Scgn* and *Srp9* in islets of NOD^k^ compared to BALB/c and B10^k^ mice were found; whereas, higher expression of *Chga* and lower expression of *Sst, Gcg, Dcx, Lmo2*, and *Mafb* were found in B10^k^ mice compared to NOD^k^ and BALB/c mice (Fig. 8e-g).

**Fig. 8.**
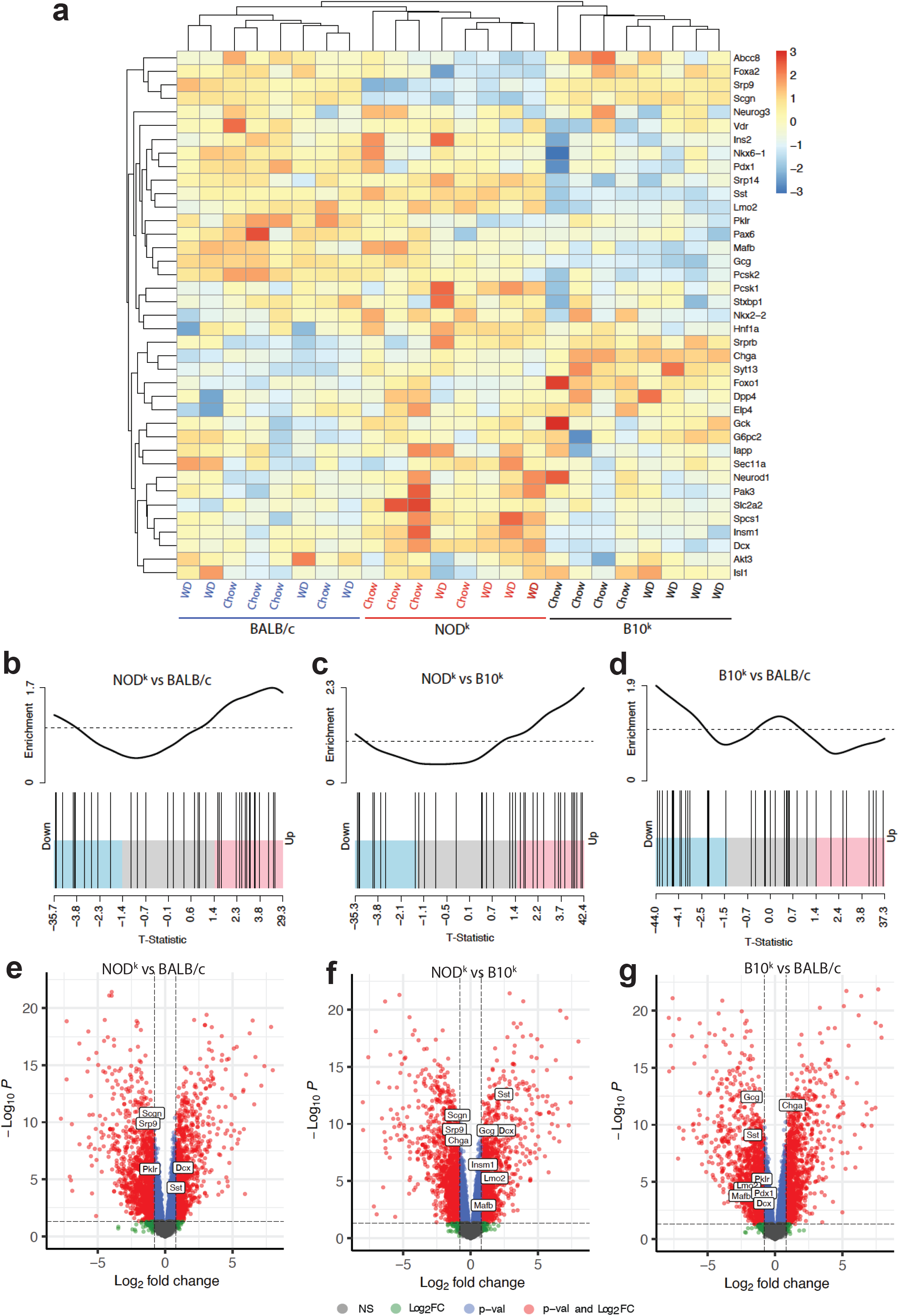
Differential expression analyses of the HALLMARK_PANCREAS_BETA_CELLS genes within bulk pancreatic islet transcriptomes of 8-week-old male BALB/c, B10^*k*^ and NOD^*k*^ mice continued either on Chow diet or in response to a 5-day WDC. **(a)** Heat map showing relative mRNA expression of the 39 genes of the HALLMARK_PANCREAS_BETA_CELL gene set. **(b-d)** Barcode plots showing enrichment of the HALLMARK_PANCREAS_BETA_CELLS genes in islets of NOD^k^ compared to BALB/c mice **(b)**, NOD^*k*^ compared to B10^*k*^ mice **(c)**, and B10^*k*^ compared to BALB/c islets **(d)**; X-axis shows T-statistic, black bars represent the 39 genes, the worm shows relative enrichment **(b-d). (e-g)** Volcano plot of DEGs between NOD^k^ and BALB/c mice **(e)**, NOD^k^ and B10^k^ mice **(f)**, and B10^k^ and BALB/c mice (g) with HALLMARK_PANCREAS_BETA_CELLS significant DEGs labelled. Dashed line significance threshold *p*<0.05, log fold-change threshold >0.8 **(e-g)**. Chow and WD-fed mice within each strain combined **(c-g)**.

Biological pathway findings derived from filtering the EMBL-EBI 60-day murine beta-cell gene expression atlas are shown in ESM Tables 1-3. Islets from NOD^*k*^ and BALB/c mice had altered between strain transcriptomic profiles of endoplasmic reticulum to golgi vesicle transport, insulin processing, intracellular protein and vesicular transport beta-cell genes (ESM Table 1). Chow-fed B10^*k*^ and BALB/c islet transcriptomes showed differences between the strains in mitochondrial function and endoplasmic reticulum/golgi pathways (ESM Table 3), with a suggestion of similar changes in mitochondrial function between NOD^*k*^ and B10^*k*^ strains, however, after false discovery adjustments, none of the NOD^k^ and B10^k^ pathway changes reached statistical significance (ESM Table 2).

## Discussion

Male NOD^*k*^ mice, through overexpression of the HEL protein in beta-cells, are prone to non-immune beta-cell failure and diabetes [16]. Here we show that male NOD^*k*^ mice simply fed a WD, rapidly develop hyperinsulinaemia, gain excess body weight and progress on to develop a type 2 diabetes phenotype, without evidence of insulitis, closely resembling that of the severe insulin resistant subtype of human type 2 diabetes. The lipid and adiponectin changes and intrapancreatic fat in NOD^*k*^ mice, even with Chow diet feeding, suggest that NOD^*k*^ mice are prone to metabolic syndrome, which could be in response to increased insulin secretion and hyperinsulinaemia [7]. In contrast, we show B10^*k*^ mice to be a hypoinsulinaemic mouse strain with already very poor glucose tolerance on Chow diet that, despite their poor insulin secretion, are resistant to developing WD-induced diabetes. Of note, B10^k^ mice were also resistant to diabetes in response to transgene-induced beta-cell stress [16]. These finding support the view that a propensity to hyperinsulinaemia increases the risk, whereas a limitation in the insulinaemic response to poor diet reduces the risk for the development of obesity-related type 2 diabetes. These findings are in keeping with human studies that show curtailing of insulin secretion in obesity can have metabolic benefit [23-26]. Preclinical studies have also shown that attenuating insulin secretion in various mouse models can prevent diet-induced obesity, insulin resistance and hyperglycaemia [27-29].

In further support of the insulin hypersecretion hypothesis, young male Chow-fed NOD^*k*^ mice display a hyperinsulinaemic hypoglycaemia phenotype, and with WD feeding for only 5 days they develop marked hyperinsulinaemia despite remaining relatively hypoglycaemic even after an intraperitoneal glucose load. The hyperinsulinaemia in Chow-fed NOD^*k*^ mice is not the result of compensation for insulin resistance, as hypoglycaemia would not be expected if this was the case, and the IPITT results showed Chow-fed NOD^*k*^ mice to be insulin sensitive. However, insulin resistance must develop rapidly with WD feeding in NOD^*k*^ mice, as the marked hyperinsulinaemia after only 5 days of WD feeding did not worsen the hypoglycaemia. Our findings in the NOD^*k*^ parallel findings in children and adults with an insulin hypersecretion phenotype [24, 30].

By 14 weeks of age, at which time glucose intolerance and very marked hyperinsulinaemia were present in WD-fed NOD^*k*^ mice, a defect in insulin clearance had developed, as very high insulin levels were discordant with nearer to normal C-peptide levels. Whether impaired insulin clearance is a primary abnormality or develops secondary to hyperinsulinaemia remains uncertain.

The *ex-vivo* insulin secretion studies did show some evidence of heightened insulin secretion in NOD^*k*^ islets, although this was not fully congruent with the marked *in-vivo* hyperinsulinaemia of these mice. Interference in the insulin assay by IAA, causing aberrant measurement was excluded. The increased percentage of proinsulin granules in mice exposed to the WD for only 5 days suggests insulin processing dysfunction, favouring a primary beta-cell defect in these mice.

The bulk islet RNAseq data of 8 week old mice show major whole islet transcriptome differences between the strains, which would again favour intrinsic hyper-responsiveness and hypo-responsiveness of islet beta-cells of, respectively, NOD^*k*^ and B10^*k*^ mice, underpinning their respective metabolic phenotypes. Of note is higher expression of *Ckb, Pgm2l1, Slco1a6*, an*d Slc39a8* in NOD^*k*^ islets compared to the other 2 strains. *CKB* (creatine kinase brain-type), important in regenerating ATP, was shown to be increased in islets from a person with hyperinsulinaemic hypoglycaemia due to an activating mutation of glucokinase [31]. PGM2L1 (phosphoglucomutase 2 like-1), also involved in phosphate transfer, is important for the synthesis of glucose-1,6-bisphospate which may regulate glycolysis in islets [32]. SLCO1A6 (Solute Carrier Organic Anion Transporter Family Member 1A6) transports bile acids like taurocholic acid (TCA), is expressed in islet beta-cells, and jejunal infusion of TCA in humans enhances glucose stimulated insulin secretion [33, 34]. *Slc39a8*, which encodes for ZiP8 a Zn^2+^ importer, had a ~10 fold increase in expression in NOD^k^ islets. ZiP8 is believed to regulate beta-cell cytosolic Zn^2+^ availability for uptake into insulin vesicles via ZnT8 [35]. Furthermore, a genetic variant of *SLC39A8* is strongly associated with type 2 diabetes and steatohepatitis [36]. Reduced islet fatty acid oxidation in islets is associated with increased insulin secretion [37, 38], such that the finding of lower expression of *Acadl* (long chain acyl-CoA dehydrogenase) and increased

*Rasgrp1* (Ras guanine nucleotide releasing protein 1) expression, a non-kinase signalling molecule activated by the fatty acid signalling molecule diacylglycerol (DAG) [39, 40], in NOD^*k*^ mice may also be relevant to the islet phenotype of this strain.

The findings of intracytoplasmic inclusion bodies in WD-fed NOD^*k*^ endocrine cells, increased endocrine cell apoptosis and changes in biological pathways of protein processing and vesicle transport suggest that NOD^*k*^ mice have dysfunctional insulin processing. This could certainly lead to loss of beta-cell mass and diabetes. Further examination of insulin processing via golgi and insulin granule trafficking in NOD^*k*^ mice is clearly warranted [41, 42].

The B10^*k*^ RNAseq data suggests a reduced islet endocrine cell differentiation (lower expression of *Sst, Gcg* (islet hormones) *Dcx, Lmo2*, and *Mafb* (islet transcription factors), and although needing to be interpreted carefully, mitochondrial metabolism pathways showed up as being altered in B10^*k*^ compared to the other strains. Defective mitochondrial metabolism deserves further investigation in this mouse model as it would impair insulin secretion.

Considering the rapidity of onset and the marked severity of the hyperinsulinaemia they develop, with onset of a non-immune diabetes by 13-24 weeks of age, the novel WD-fed NOD^*k*^ mice appears to be highly suitable as a murine model of the severe insulin resistant subtype of type 2 diabetes. Obvious advantages compared to other mice models of type 2 diabetes are that it breeds well, it is not dependent on genotyping or monogenetic obesity, chow-fed mice remain healthy, and a second hit such as streptozotocin or alloxan is not needed [14, 43, 44].

Limitations in this work include that female mice were minimally studied due to their resistance to developing diabetes. IPITTs were performed in Chow fed mice only at 13 weeks of age, as widely disparate fasting glucose levels in WD-fed mice would make interpretation difficult, and hyperinsulinaemic-euglycaemic insulin clamps were not performed. Performing more detailed assessments of insulin resistance however would likely not to be helpful, as insulin resistance and hyperinsulinaemia almost always accompany each other.

In conclusion, male WD-fed NOD^*k*^ mice exhibit a striking predisposition to hyperinsulinaemia that progresses to overt diabetes, resembling the severe insulin resistant subtype of type 2 diabetes. In contrast, B10^*k*^ mice, despite a hypoinsulinaemic phenotype with poor glucose tolerance on Chow diet, they are resistant to WD-induced diabetes. While the underlying mechanisms are likely multifactorial and require further investigation, these findings are consistent with the insulin hypersecretion hypothesis [6, 7, 9, 10], and the concept of vulnerability of hypersecreting beta-cells [45]. Importantly, the WD-fed NOD^k^ mouse emerges as a highly promising new model for investigating the pathogenesis of severe insulin resistant type 2 diabetes and evaluating potential therapeutic targets.

## Supporting information

Electronic Supplementary Material

## Abbreviations

*ANU*: Australian National University
*BALB/c*: Bagg Albino
*B10^k^*: B10.BR-*H2*^*k*^/SgSnJ
*Chow*: Standard rodent chow diet
*DEGs*: Differentially expressed genes
*EM*: Electron microscopy
*HEL*: Hen egg lysozyme
*IBMX*: 3-isobutyl-1-methylxanthine
*NOD*: *Non-obese diabetic*
*NOD*^*k*^: NOD.BR-*H2*^*k*^/Wicker
*PEG*: Polyethylene glycol
*WD*: Western diet
*WDC*: Western diet challenge

## Acknowledgments

We acknowledge the technical assistance of Ms Elaine Bean in preparation of tissue sections for histological analysis and Ms Sabine Grüninger for the preparation of the electron microscopy samples, as well as the staff of The Canberra Hospital Animal Facility, the Australian Phenomics Facility (ANU), and the Biomolecular Resource Facility (ANU). We thank Prof Marc Prentki (University of Montreal) for his generous and constructive discussions.

## Data availability

The data from the bulk islet RNA-seq is deposited in GEO database (accession no. XXXX). All the other data associated with this manuscript can be found in the manuscript, the ESM, or can be requested from the corresponding author (CJN).

## Contribution statement

MFW, AH, VD-A and CJN conceived and designed the study. MFW, AKH, VD-A, MS and TD performed experiments and analysed data. MEK and VD-A analysed electron microscopy and JED analysed histological data. AB, Z-PF and TDA performed analyses of the RNAseq data. MFW and CJN wrote the manuscript. All authors provided critical revisions and edits to the manuscript. All authors read and approved the final manuscript. CJN is the guarantor for this study.

