## Supplementary material for "A propensity to hyperinsulinaemia underpins diet-induced diabetes in NOD^*k*^ mice": Electronic Supplementary Material

Waters MF et al.

**Electronic supplemental material (ESM)**

ESM Materials and Methods	Pages 1-4

ESM Fig. 1	Female NOD*^k^* fed glucose and IPGTT	Page 5

ESM Fig. 2	IAA assessment	Page 6

ESM Fig. 3	Pancreas histology H&E 14 weeks of age	Page 7

ESM Fig. 4	Pancreas insulin immunohistochemistry	Page 8

ESM Fig. 5	Bulk RNAseq top 50 DEGs	Page 9

ESM Fig. 6	Islet mRNA expression by qRT-PCR	Page 10

ESM Table 1	NOD*^k^* vs BALB/c ENRICHR pathway analysis	Page 11

ESM Table 2	NOD*^k^* vs B10*^k^* ENRICHR pathway analysis	Page 12

ESM Table 3	B10*^k^* vs BALB/c	ENRICHR pathway analysis	Page 13

**ESM Materials and Methods**

**Intraperitoneal glucose tolerance and insulin tolerance testing** ipGTTs were performed in conscious mice after fasting for 6 hours (08:00 until 14:00). Tail vein fasting blood glucose (glucose meter) and a blood sample (30 μl) were collected for plasma insulin (time 0 min), followed by the administration of 1g/kg 25% glucose solution via intraperitoneal (i.p.) injection. Tail vein blood glucose was then measured at 5, 15, 30, 60, 90, 120 min and additional blood samples (30 μl) for plasma insulin were taken at 15 and 90 min. Area under the curve (AUC) for i.p.GTT glucose and insulin used the trapezoid method. Insulin tolerance tests (ipITTs) were performed in conscious mice after fasting for 4 hours (0:800 until 12:00). After a time 0 min tail blood glucose, 0.75U regular insulin (Humulin R, Eli Lilly), diluted 0.1 U/ml in sterile normal saline, was administered i.p. and tail blood glucose was then measured at 15, 30, 45, 60 and 90 min.

**Insulin autoantibodies (IAA) assessment**

Presence/interference of IAA was tested using two methodologies, the non-competitive ELISA method as previously described and optimised [1, 2], and the Polyethylene glycol (PEG) precipitation method [3].

The non-competitive ELISA method was performed in a 96 well ELISA plate (Cat# 423501, BioLegend, San Diego, CA, USA) coated overnight at 4°C with 100 μL of coating buffer (Cat# 421701, BioLegend) with or without 50 μg/mL human insulin (Actrapid, Novo Nordisk, Denmark). After washing 3 times (wash buffer, 50 mM Trizma hydrochloride and 0.2% Tween 20 in milli-Q water, Merck Sigma-Aldrich), the wells were blocked with 100 μl PBS containing 2% BSA (cat# A7030, Merck Sigma-Aldrich) for 2 h at room temperature (RT). After blocking buffer removal, 5 μl of mice plasma samples diluted 1:10 with 45 μl assay buffer (PBS containing 2% BSA) or 50 μl assay buffer as sample blank, all in duplicate, were added to wells at RT for 4 h, followed by washing x4, then incubation with biotinylated anti-mouse IgG1 (1:10,000) (Cat# ab11587, Abcam, Cambridge, UK) for 30 min at RT, followed by washing x4, then incubation with avidin-horseradish peroxidase (1:1,000) (Cat# 405103, BioLegend) for 15-20 min at RT, followed by washing x5, and finally incubation with 100 μl of 3,3', 5,5"-tetramethylbenzidine solution (Cat# 421101, BioLegend) for 10-15 min at RT. The reaction was stopped by adding 100 µL of stop solution (1N H_2_SO_4_) into each well. Starting with the blocking step, all incubations were on a rocker (60 rpm). Absorbance was then measured at 450 nm using an Infinite 200 PRO plate reader (Tecan Trading AG, Switzerland). The absorbance of test samples without plate-bound insulin (background) were subtracted from absorbance of test samples with plate-bound insulin for actual sample absorbance calculation.

For the PEG method, plasma insulin levels were measured in duplicate samples prepared with or without PEG precipitation. For PEG precipitation samples (for IAA bound insulin clearance), 10 μL of plasma samples were initially diluted with 15 μL of the ELISA kit diluent, then mixed with 25μl of 25% w/v PEG 6000 (Cat# 81260, Merck Sigma-Aldrich) diluted with ELISA kit diluent. For the samples without PEG precipitation, 10 μl of plasma samples were diluted with 40 μl of the ELISA kit diluent. Samples were then vortexed for 10 s, allowed to stand on ice for 5 min before centrifugation (10, 000*g*) for 15 min at RT. The supernatants were then collected and used for the determination of insulin concentration using the ultra-sensitive mouse insulin ELISA kit (Cat #90080, Crystal Chem, Illinois, USA).

**Pancreas insulin immunohistochemistry**

From the pancreas paraffin blocks, three consecutive sections of 5μm at distance of 100 μm were cut and collected on HDS microscope slides (HD Scientific Supplies, Kings Park, NSW, Australia). The slides were de-paraffinised through two washes in xylene solvent and rehydrated in graded ethanol solutions (100, 90 and 70%). The slides were then washed in cold water and immersed in PBS (PBS; Cat# 09-8912-100, Medicago AB, Uppsala, Sweden) to prevent drying. To reduce background or unspecific staining, slides were blocked in 5% fetal bovine serum (Hi FBS; Cat# 10082-147, Gibco, Life Technologies Corp., NY, USA) solution for 15 minutes. The slides were then incubated for 1 hour with a primary antibody, polyclonal guinea-pig anti-insulin (diluted 1:100 in 5% FBS/PBS solution) (Cat# A0564, Dako, CA, USA), in a humidified incubation chamber at room temperature. Unbound primary antibody was removed by washing twice in PBS. Endogenous peroxidase was blocked by 20 minutes incubation with 3% hydrogen peroxide (H_2_O_2_) made with 1x PBS solution. Following another wash (2x) in PBS, the slides were stained with a secondary antibody, polyclonal rabbit anti-guinea pig horseradish peroxidase (diluted 1:40 in 5% FBS/PBS solution) (Cat# P0141, Dako, Glostrup, Denmark) for 45 minutes in a humidified incubation chamber at room temperature. Unbound secondary antibody was washed out with PBS (2 minutes, 2x) and slides were stained with 450 μL of 3,3’-Diaminobenzidine (DAB) (DAB; Cat# D4293, SIGMAFAST^™^ Sigma-Aldrich, Castle Hill, NSW, Australia) for exactly 2 minutes for colour development. The slides were counterstained with Harris Haemotoxylin solution (Cat #HHX2.5L, POCD, Artarmon, NSW, Australia) for 2 minutes and rinsed under running water. They were then immersed in 1% sodium bicarbonate solution (Cat# S3817, Sigma-Aldrich CA, USA) for 15 seconds and again washed under tap water. Finally, the slides were dehydrated in graded ethanol solutions (70, 90 and 100%), and incubated in xylene solvent through two washes (2 minutes each). The slides were then cover-slipped using a small amount of Pertex mounting medium (Cat# 41-4012-00 MEDITE Medical GmbH, Burgdorf, Germany).

**Beta-cell mass measurement**

Images of pancreas sections immunostained for insulin were obtained using an Aperio slide scanner (Aperio ScanScope XT, **Aperio Technologies, Inc, CA, USA) at 20x magnification. The images were opened in Image J 1.48v software (Wayne Rasband, National Institute of Health, USA) and cleaned of debris, fat or lymph node tissue by using the software. Imaging** threshold adjustments were set for selection of total pancreas area and the areas immunostained for insulin, with Image J measurements of both being recorded on an excel spread sheet. This process was repeated for each of the 3 levels. For each level, the percentage β-cell area per pancreas was calculated (islet sum area x 100 / pancreas area). Percentage areas of the 3 sections were then averaged. Finally, the β-cell mass was calculated, when the percentage β-cell area per pancreas was multiplied by the pancreatic weight (mg).

**Islet ex-vivo insulin secretion**

To assess ex vivo insulin secretion, freshly isolated islets were incubated overnight in RPMI-1640 medium supplemented with 10% fetal calf serum, 10 mmol/l HEPES (pH 7.4), 1 mmol/l sodium pyruvate, 100 U/ml penicillin and 100 µg/ml streptomycin (RPMI complete medium) with 11 mmol/l glucose at 37°C in a humidified atmosphere containing 5% CO_2_. The next morning, batches of 8 islets in triplicate (technical repeats for each data point) were distributed into 24 well microplates and incubated for 2 hours in RPMI complete supplemented with 3 mmol/l glucose and then pre-incubated in Krebs Ringer Bicarbonate buffer with 10 mmol/l HEPES at pH 7.4 and 0.5% BSA (KRBH-BSA buffer), containing 3 mM glucose, for 40 minutes at 37 degrees with 5% CO_2_. The islets were then washed in KRBH-BSA buffer and incubated in 1.2 ml/well KRBH-BSA buffer with 3 mmol/l glucose (3G) and 1 ml/well of KRBH-BSA buffer with either 8 mmol/l glucose (8G), 16 mM glucose (16G), or 8 mM glucose with 100 μM isobutylmethylxanthine (IBMX, Sigma-Aldrich, Bayswater, VIC, Australia) (8G+IBMX) for 45 mins at 37^0^C. An aliquot of 200 uL of incubation solution was taken from wells with 3 mM glucose at 5 minutes to determine the background insulin concentration. After 45 minutes the incubation solution was collected to determine secreted insulin using the insulin ultrasensitive kit (Cisbio, Codolet, France) and islets were transferred to tubes containing 75% ethanol acidified with 1.5% hydrochloric acid. Islet intracellular insulin content was later released by sonication (Omni Ruptor 250, Omni International, GA, USA) and measured using the insulin ultrasensitive kit.

**Islet electron microscopy**

Freshly isolated islets were washed in PBS and fixed in 2 % glutaraldehyde in 0.1 M cacodylate buffer (Electron Microscopy Sciences, Morgantown, PA, USA), pH 7.4, overnight at 4°C. After consecutive washes in 0.1 M cacodylate buffer, the islet pellets were postfixed in 4 % osmium in the cacodylate buffer (Electron Microscopy Sciences, USA) for 1 h at room temperature. Two washes in distilled water were followed by block staining with 2 % aqueous uranyl acetate (ProSciTech, Thuringowa Central, QLD, Australia). The islets were then dehydrated through a series of ethanol and embedded in TAAB epoxy resin (TAAB Laboratories Equipment Ltd, Berks, UK. Ultrathin sections (90 nm) were cut, mounted on 150-mesh copper/palladium grids and stained with Reynolds lead citrate. Representative digital electron micrographs of individual beta cells were captured from 2-3 beta-cells per islet from 5-6 different islets of each mouse (giving 10-12 beta-cell images per mouse) by transmission electron microscopy (JEOL 1011; JEOL Ltd, Tokyo, Japan) using a digital camera (MegaView G2; Olympus, Münster, Germany). Images were analyzed with iTEM software v.5.2 (Olympus, Münster, Germany). Mature and immature insulin granules were identified based on core density and presence or absence of the characteristic mature granule halo. The assessments were performed blinded.

[1]	Jhala G, Chee J, Trivedi PM, et al. (2016) Perinatal tolerance to proinsulin is sufficient to prevent autoimmune diabetes. JCI Insight 1(10): e86065. 10.1172/jci.insight.86065

[2]	Babaya N, Liu E, Miao D, Li M, Yu L, Eisenbarth GS (2009) Murine high specificity/sensitivity competitive europium insulin autoantibody assay. Diabetes Technol Ther 11(4): 227-233. 10.1089/dia.2008.0072

[3]	Huynh T (2020) Clinical and Laboratory Aspects of Insulin Autoantibody-Mediated Glycaemic Dysregulation and Hyperinsulinaemic Hypoglycaemia: Insulin Autoimmune Syndrome and Exogenous Insulin Antibody Syndrome. Clin Biochem Rev 41(3): 93-102. 10.33176/AACB-20-00008

**ESM Figures**

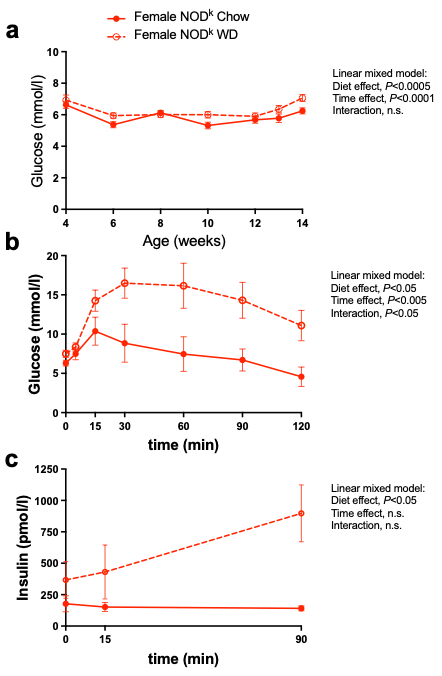

**ESM Fig. 1.** Fed-state blood glucose and intraperitoneal glucose tolerance tests in female NOD*^k^* mice fed Chow or western diet (WD) from 4 to 14 weeks of age. **(a)** 9am fed blood glucose; *n*=11 Chow diet, *n*=12 WD. **(b-c)** Intraperitoneal glucose tolerance tests (ipGTTs) performed at 13 weeks of age; blood glucose, *n*=8 both groups **(b)**, Plasma insulin, *n*=7 both groups **(c)**. Data are shown as means ± SEM. Analysed with repeated measures two way ANOVA or mixed model, as indicated.

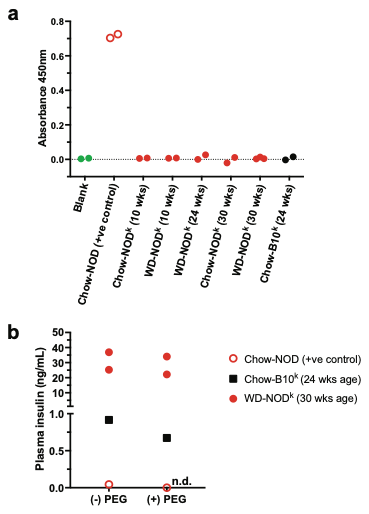

**ESM Fig. 2.** Insulin autoantibody assessment. **(a)** Using a non-competitive ELISA method and (**b)** a plasma insulin assay after polyethylene glycol (PEG) precipitation. Diabetic male Chow-fed NOD mice (positive control) and male Chow and WD-fed NOD*^k^* mice at 10, 24 and 30 weeks of age and Chow-fed B10*^k^* mice at 24 weeks of age were tested. Each data point is from the average of duplicate samples.

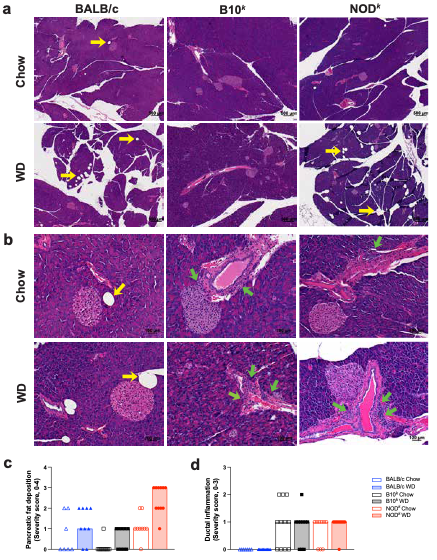

**ESM Fig. 3.** Pancreas histology in BALB/c, B10*^k^* and NOD^k^ mice fed Chow and WD at 14 weeks of age. **(a)** Representative low power images of pancreatic sections stained with H&E showing presence of fat deposits; scale bar 500 μm, initial magnification x50; yellow arrows point to fat deposits. **(b)** Representative high power images of pancreas sections stained with H&E showing periductal inflammation; scale bar 50 μm, initial magnification x400; green arrows point to areas of inflammation. **(c, d)** Pancreatic fat deposition scores; score matrix 0-none, 1-minimal, 2-mild, 3-moderate, 4-severe **(c)**, Periductal inflammation scores; score matrix 0-none, 1-<10%, 2-10-50%, 3->50% of ducts affected **(d)**; *n*=7 BALB/c Chow, *n*=10 BALB/c WD, *n*=11 B10*^k^* Chow, *n*=9 B10*^k^* WD, *n*=9 NOD*^k^* Chow, *n*=11 NOD*^k^* WD. **(c, d).** Each data point represents one mouse and the bars are the group median. Ordinal logistic regression model: diet effect, *p*=<0.0001, strain effect, *p*=<0.0001, diet×strain effect, *p*=not significant **(c)**; Kruskal-Wallis rank sum test: diet effect, *p*=0.95, strain effect, *p*<0.0001 **(d)**.

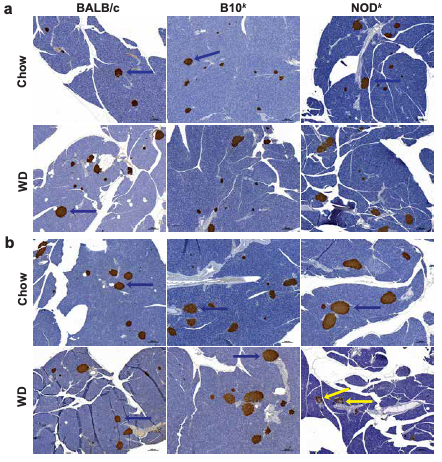

**ESM Fig. 4.** Pancreas insulin immunohistochemistry in BALB/c, B10*^k^* and NOD*^k^* mice fed Chow and WD. **(a)** Representative low power images of pancreatic sections from mice at 14 weeks of age immunostained for insulin. **(b)** Representative low power images of pancreatic sections from mice at 24 weeks of age, or at time of harvest if diagnosed with diabetes, immunostained for insulin. Insulin staining brown - shown by red arrows. Yellow arrows point to islets with patchy and less intense insulin staining. Scale bar 500 μm, initial magnification x50.

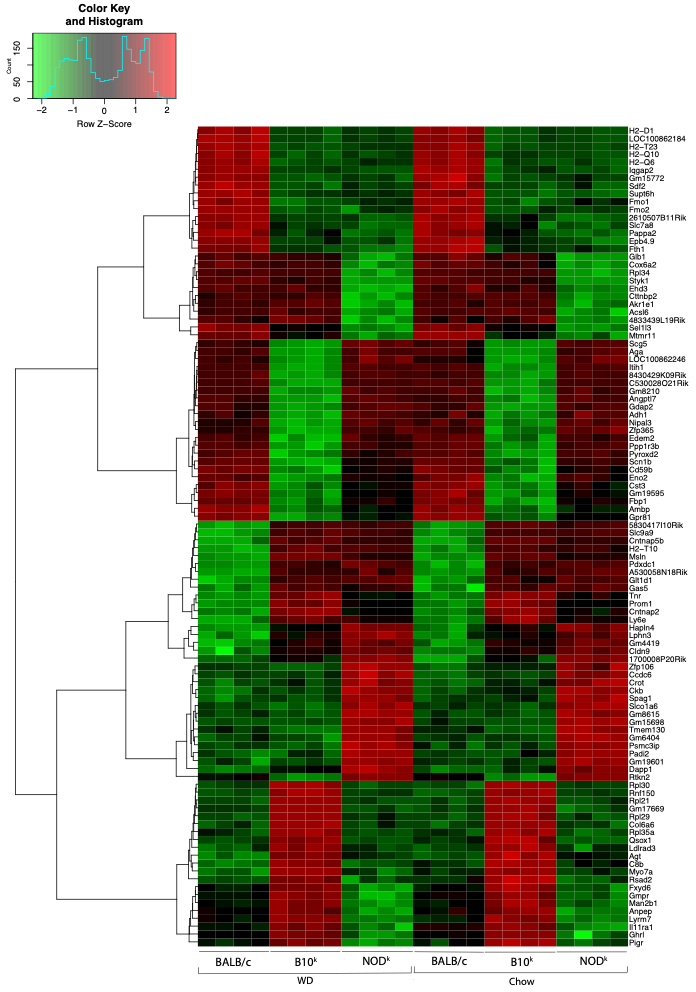

**ESM Fig. 5.** Heatmap of top differentially expressed genes from bulk islet RNAseq from 8 week old BALB/c, B10*^k^* and NOD*^k^* mice continued on Chow diet or in response to a 5 day WD challenge.

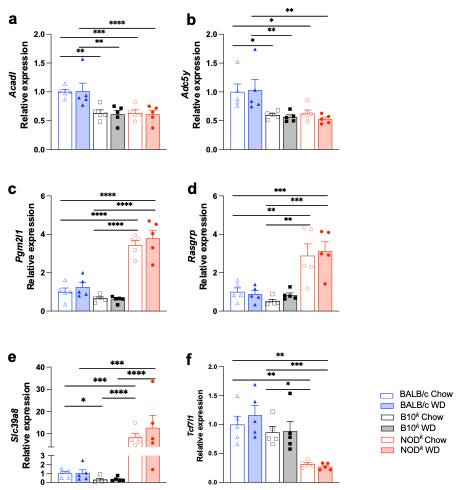

**ESM Fig. 6.** Islet mRNA expression by qRT-PCR for validation of RNAseq in genes of interest from 8 week old BALB/c, B10*^k^* and NOD*^k^* mice continued either on Chow diet or in response to a 5-day WD challenge. (**a**) *Acadl*, (**b**) *Adc5y*, (**c**) *Pgm2l1*, (**d**) *Rasgrp*, (**e**) *slc39a8*, (**f**) *Tcf7l1*. Data are shown as means ± SEM; *n* = 5. Data analysed by two-way ANOVA, with Tukey’s multiple comparison *post-hoc* tests **(c, d)**, **p*<0.05, ***p*<0.01, *****p*<0.0001.

**ESM Table 1.** **Top 10 beta-cell specific biological processes for comparisons according to Chow diet or WD of NOD*^k^* compared to BALB/c on ENRICHR pathway analysis**

| **Pathway Term** | **Overlap** | **P-value** | **Adjusted P-value** | **Odds Ratio** | **Genes** |
| --- | --- | --- | --- | --- | --- |
| **NOD^I^ Chow vs BALB/c Chow** |  |  |  |  |  |
| Endoplasmic Reticulum To Golgi Vesicle-Mediated Transport (GO:0006888) | 19/115 | 2.22E-12 | 2.22E-09 | 9.87 | RAB2A;COPB2;ARF4;RAB1A;TRAPPC2L;TMED10;SAR1B;COPB1;YIPF5;MIA3;TRAPPC9;ARCN1;LMAN1;SCFD1;TMED2;KDELR2;TRAPPC13;KDELR3;BET1 |
| Golgi Vesicle Transport (GO:0048193) | 24/197 | 2.64E-12 | 2.22E-09 | 6.98 | RAB2A;ARF4;COPB2;RAB1A;TRAPPC2L;TMED10;COG6;SAR1B;COPB1;YIPF5;COPZ2;MIA3;TRAPPC9;SORL1;ARCN1;LMAN1;SCFD1;EHD3;TMED2;KDELR2;KDELR3;TRAPPC13;PLPP3;BET1 |
| Retrograde Vesicle-Mediated Transport, Golgi To Endoplasmic Reticulum (GO:0006890) | 8/48 | 5.36E-06 | 0.0030 | 9.73 | ARF4;COPB2;SCFD1;TMED10;COPZ2;KDELR2;KDELR3;PLPP3 |
| Vesicle-Mediated Transport (GO:0016192) | 22/411 | 4.47E-05 | 0.0188 | 2.80 | RAB2A;TBC1D8B;ARF4;RAB1A;SYT4;TMED10;VPS4B;GDI2;ARL1;RAB39B;MIA3;VAMP8;RAB10;AP3M1;EHD3;DNER;KIF5A;OCIAD1;TMED2;ATP6V1H;PIK3C3;RAB5A |
| Insulin Processing (GO:0030070) | 3/5 | 8.29E-05 | 0.0279 | 72.19 | PCSK2;YIPF5;P4HB |
| Endoplasmic Reticulum Organization (GO:0007029) | 8/54 | 1.12E-04 | 0.0289 | 7.22 | RAB10;REEP1;TRAM1;REEP5;CCDC47;MIA3;RTN4 |
| Endoplasmic Reticulum Organization (GO:0007029) | 7/54 | 1.12E-04 | 0.0289 | 7.22 | ERP44;VCP;UFM1;TMEM117;CCDC47;DNAJB9;TRIB3;ANKS4B;P4HB;HSP90B1 |
| Vesicle Coating (GO:0006901) | 5/28 | 2.35E-04 | 0.0474 | 10.50 | TMED10;TRAPPC2L;TMED2;TRAPPC13;TRAPPC9 |
| Vacuolar Acidification (GO:0007035) | 5/29 | 2.80E-04 | 0.0474 | 10.06 | ATP6V0B;CCDC115;ATP6V1H;ATP6V0E2;ATP6V0C |
| Aromatic Amino Acid Transport (GO:0015801) | 3/7 | 2.82E-04 | 0.0474 | 36.09 | SLC25A29;SLC7A8;SLC3A2 |
| **NOD*^k^* WD vs BALB/c WD** |  |  |  |  |  |
| Golgi Vesicle Transport (GO:0048193) | 16/197 | 1.60E-06 | 0.0023 | 4.62 | COPB2;RAB1A;TMED9;TRAPPC2L;TMED10;SAR1B;COPZ2;MIA3;SORL1;LMAN1;EHD3;RABIF;VAPB;TMED3;KDELR3;PLPP3 |
| Endoplasmic Reticulum To Golgi Vesicle-Mediated Transport (GO:0006888) | 12/115 | 2.45E-06 | 0.0023 | 6.04 | COPB2;RAB1A;LMAN1;SEC13;TMED9;TRAPPC2L;TMED10;SAR1B;VAPB;TMED3;MIA3;KDELR3 |
| Cellular Response To Amino Acid Starvation (GO:0034198) | 7/42 | 1.47E-05 | 0.0093 | 10.28 | DAP;RRAGA;DAPL1;RRAGD;EIF2AK3;ITFG2;MAPK3 |
| COPII-coated Vesicle Budding (GO:0090114) | 6/36 | 6.10E-05 | 0.0271 | 10.25 | RAB1A;SEC13;TMED10;TRAPPC2L;VAPB;MIA3 |
| Intracellular Protein Transport (GO:0006886) | 18/325 | 7.15E-05 | 0.0271 | 3.06 | CD74;COPB2;RAB1A;SEC13;TMED9;TMED10;SAR1B;ARL1;COPZ2;XPO7;SORL1;NUP93;LMAN1;TMED3;SCG5;IFT27;CALR;RAB5A |
| Response To Amino Acid Starvation (GO:1990928) | 6/41 | 1.30E-04 | 0.0411 | 8.80 | DAP;RRAGA;DAPL1;EIF2AK3;ITFG2;MAPK3 |
| Cellular Response To Starvation (GO:0009267) | 10/126 | 1.76E-04 | 0.0477 | 4.45 | GABARAPL1;RRAGA;DAP;DAPL1;RRAGD;EIF2AK3;PCSK9;PIK3C3;ITFG2;MAPK3 |
| Aromatic Amino Acid Transport (GO:0015801) | 3/7 | 2.39E-04 | 0.0568 | 38.20 | SLC25A29;SLC7A8;SLC3A2 |
| Axonogenesis (GO:0007409) | 12/188 | 3.19E-04 | 0.0643 | 3.52 | LGI1;NELL1;KIF5C;SLITRK1;FEZ1;ATL1;PLPPR4;KIF5A;SLITRK6;TRIM46;EMB;NEFH |
| Lysosomal Lumen Acidification (GO:0007042) | 4/18 | 3.44E-04 | 0.0643 | 14.58 | ATP6V0B;CCDC115;ATP6V1H;ATP6V0C |

**ESM Table 2.** **Top 10 beta-cell specific biological processes for comparisons according to Chow diet or WD of NOD*^k^* compared to B10*^k^* on ENRICHR pathway analysis**

| **Pathway Term** | **Overlap** | **P-value** | **Adjusted P-value** | **Odds Ratio** | **Genes** |
| --- | --- | --- | --- | --- | --- |
| **NOD*^k^* Chow vs B10*^k^* Chow** |  |  |  |  |  |
| Mitochondrial ATP Synthesis Coupled Electron Transport (GO:0042775) | 7/70 | 1.18E-04 | 0.176 | 7.06 | NDUFA12;NDUFS5;NDUFA3;NDUFS2;SDHA;COX6A2;UQCRH |
| Protein Targeting To ER (GO:0045047) | 4/23 | 4.27E-04 | 0.236 | 13.27 | SPCS3;SPCS1;SRP14;SRP9 |
| Oxidative Phosphorylation (GO:0006119) | 6/63 | 4.73E-04 | 0.236 | 6.66 | NDUFA12;NDUFS5;NDUFA3;NDUFS2;SDHA;UQCRH |
| Aerobic Electron Transport Chain (GO:0019646) | 6/68 | 7.13E-04 | 0.266 | 6.13 | NDUFS5;NDUFA3;NDUFS2;SDHA;COX6A2;UQCRH |
| Proton Motive Force-Driven Mitochondrial ATP Synthesis (GO:0042776) | 5/53 | 1.47E-03 | 0.306 | 6.58 | NDUFA12;NDUFS5;NDUFA3;NDUFS2;SDHA |
| Establishment Of Protein Localization To Endoplasmic Reticulum (GO:0072599) | 3/16 | 0.0019 | 0.306 | 14.50 | SPCS3;SPCS1;SRP14 |
| Response To Endoplasmic Reticulum Stress (GO:0034976) | 7/114 | 0.00229 | 0.306 | 4.14 | ERN1;ERLEC1;VCP;CREB3L1;TMEM117;RASGRF2;CREB3L2 |
| Cellular Respiration (GO:0045333) | 6/85 | 0.0023 | 0.306 | 4.80 | NDUFA12;NDUFS5;NDUFA3;NDUFS2;SDHA;UQCRH |
| Aerobic Respiration (GO:0009060) | 5/59 | 0.0024 | 0.306 | 5.84 | NDUFA12;NDUFS5;NDUFA3;NDUFS2;UQCRH |
| Regulation Of Relaxation Of Cardiac Muscle (GO:1901897) | 2/5 | 0.0024 | 0.306 | 41.78 | CHGA;PDE4B |
| **NOD*^k^* WD vs B10*^k^* WD** |  |  |  |  |  |
| Antigen Processing And Presentation Of Peptide Antigen Via MHC Class I (GO:0002474) | 5/18 | 4.57E-05 | 0.0665 | 16.21 | PDIA3;SAR1B;TAP2;CALR;B2M |
| Regulation Of Macroautophagy (GO:0016241) | 11/96 | 8.36E-05 | 0.0665 | 4.94 | ERN1;ATP6V0B;PINK1;EXOC4;UBQLN1;QSOX1;PIK3C3;ATP6V1E1;ATP6V0E2;ATP6V1D |
| Regulation Of Lysosome Organization (GO:1905671) | 3/5 | 1.23E-04 | 0.0665 | 63.00 | SCARB2;LAPTM4B;MCOLN1 |
| Protein Transport (GO:0015031) | 19/313 | 1.58E-04 | 0.0665 | 2.77 | DNAJC19;NGFR;CD74;SEC13;TMED9;COPA;SAR1B;VPS33A;ARL1;MIA2;MIA3;MCOLN1;RAB24;TMED3;SCG5;SIL1;RAB6B;MON1A;TMED4 |
| Cotranslational Protein Targeting To Membrane (GO:0006613) | 4/13 | 1.79E-04 | 0.0665 | 18.70 | TRAM1;SRP14;SIL1;SRP9 |
| Mitochondrial ATP Synthesis Coupled Electron Transport (GO:0042775) | 8/70 | 2.24E-04 | 0.0665 | 5.46 | NDUFA12;NDUFS5;NDUFA3;NDUFA1;NDUFS2;SDHA;COX6A2;UQCRH |
| Golgi Vesicle Transport (GO:0048193) | 14/197 | 2.34E-04 | 0.0665 | 3.26 | TMED9;COPA;SAR1B;PEF1;MIA2;MIA3;SORL1;EHD3;RABIF;VAPB;TMED3;PLPP3;RAB6B;TMED4 |
| Response To Endoplasmic Reticulum Stress (GO:0034976) | 10/114 | 3.47E-04 | 0.08225 | 4.08 | PDIA3;ERN1;ERLEC1;VCP;CREB3L1;TMEM117;RASGRF2;UBQLN1;P4HB;PDIA6 |
| Endoplasmic Reticulum To Golgi Vesicle-Mediated Transport (GO:0006888) | 10/115 | 3.73E-04 | 0.08225 | 4.04 | COPA;SEC13;TMED9;PEF1;SAR1B;VAPB;TMED3;MIA2;MIA3;TMED4 |
| Regulation Of Oxidative Stress-Induced Cell Death (GO:1903201) | 3/7 | 4.15E-04 | 0.0825 | 31.50 | PINK1;UBQLN1;P4HB |

**ESM Table 3.** **Top 10 beta-cell specific biological processes for comparisons according to Chow diet or WD of B10*^k^* compared to BALB/c on ENRICHR pathway analysis**

| **Pathway Term** | **Overlap** | **P-value** | **Adjusted P-value** | **Odds Ratio** |  |
| --- | --- | --- | --- | --- | --- |
| **B10*^k^* Chow vs BALB/c Chow** |  |  |  |  |  |
| Aerobic Electron Transport Chain (GO:0019646) | 10/68 | 3.13E-06 | 0.0039 | 7.52 | COX8A;NDUFA3;NDUFA1;NDUFS2;NDUFV3;SDHA;COX6A2;COX5A;UQCRH;COX6B1 |
| Mitochondrial ATP Synthesis Coupled Electron Transport (GO:0042775) | 10/70 | 4.09E-06 | 0.0039 | 7.26 | COX8A;NDUFA3;NDUFA1;NDUFS2;NDUFV3;SDHA;COX6A2;COX5A;UQCRH;COX6B1 |
| Retrograde Vesicle-Mediated Transport, Golgi To Endoplasmic Reticulum (GO:0006890) | 8/48 | 1.18E-05 | 0.0075 | 8.69 | COPA;NBAS;TMED10;TMEM115;LMAN2;COPZ2;KDELR3;PLPP3 |
| Cellular Respiration (GO:0045333) | 10/85 | 2.38E-05 | 0.0113 | 5.81 | COX8A;CHCHD5;NDUFA3;NDUFA1;NDUFS2;NDUFV3;SDHA;COX5A;UQCRH;COX6B1 |
| Golgi Lumen Acidification (GO:0061795) | 4/9 | 3.09E-05 | 0.0118 | 34.50 | ATP6V1H;ATP6V0C;ATP6V1D;ATP6V1F |
| Endosomal Lumen Acidification (GO:0048388) | 4/11 | 7.82E-05 | 0.0249 | 24.64 | ATP6V1H;ATP6V0C;ATP6V1D;ATP6V1F |
| Golgi Vesicle Transport (GO:0048193) | 14/197 | 1.84E-04 | 0.0501 | 3.34 | RAB1A;NBAS;COPA;TRAPPC2L;TMED10;COG6;SAR1B;COPZ2;LMAN1;TMEM115;LMAN2;TMED2;KDELR3;PLPP3 |
| Regulation Of Lysosomal Lumen pH (GO:0035751) | 5/25 | 2.22E-04 | 0.0530 | 10.80 | VPS33A;ATP6V1H;ATP6V1D;ATP6V0C;ATP6V1F |
| Mitochondrial Electron Transport, Cytochrome C To Oxygen (GO:0006123) | 4/16 | 3.94E-04 | 0.0784 | 14.37 | COX8A;COX5A;COX6A2;COX6B1 |
| Energy Derivation By Oxidation Of Organic Compounds (GO:0015980) | 6/43 | 4.10E-04 | 0.0784 | 7.01 | COX8A;CHCHD5;GFPT1;COX5A;UQCRH;COX6B1 |
| **B10*^k^* WD vs BALB/c WD** |  |  |  |  |  |
| Regulation Of Cardiac Muscle Hypertrophy (GO:0010611) | 4/23 | 8.37E-04 | 0.443 | 11.03 | RGS2;PRKCA;IL6ST;LMCD1 |
| Endoplasmic Reticulum Membrane Organization (GO:0090158) | 3/11 | 9.87E-04 | 0.443 | 19.61 | REEP5;ATL1;RTN4 |
| Relaxation Of Cardiac Muscle (GO:0055119) | 3/12 | 0.0013 | 0.443 | 17.43 | CHGA;GSTM2;RGS2 |
| Cellular Response To Acid Chemical (GO:0071229) | 4/27 | 0.0016 | 0.443 | 9.11 | RRAGD;LAMTOR2;LAMTOR3;KLF2 |
| Prostanoid Biosynthetic Process (GO:0046457) | 3/14 | 0.0021 | 0.443 | 14.26 | CD74;EDN2;PTGES |
| Endoplasmic Reticulum Tubular Network Organization (GO:0071786) | 3/16 | 0.0031 | 0.443 | 12.07 | REEP1;ATL1;RTN4 |
| Establishment Of Protein Localization To Endoplasmic Reticulum (GO:0072599) | 3/16 | 0.0031 | 0.443 | 12.07 | SPCS3;SPCS2;SRPRB |
| Pteridine-Containing Compound Metabolic Process (GO:0042558) | 2/5 | 0.0034 | 0.443 | 34.79 | GCH1;MTHFD1L |
| Prostaglandin Biosynthetic Process (GO:0001516) | 3/17 | 0.0037 | 0.443 | 11.20 | CD74;EDN2;PTGES |
| Anterograde Dendritic Transport (GO:0098937) | 2/6 | 0.0050 | 0.443 | 26.09 | KIF5C;KIF5A |
